# Inhibition of RNA Polymerase I Transcription Activates Targeted DNA Damage Response and Enhances the Efficacy of PARP Inhibitors in High-Grade Serous Ovarian Cancer

**DOI:** 10.1101/621623

**Authors:** Elaine Sanij, Katherine M. Hannan, Shunfei Yan, Jiachen Xuan, Jessica E. Ahern, Keefe T. Chan, Jinbae Son, Olga Kondrashova, Elizabeth Lieschke, Matthew J. Wakefield, Anna S. Trigos, Daniel Frank, Sarah Ellis, Carleen Cullinane, Jian Kang, Gretchen Poortinga, Purba Nag, Kum Kum Khanna, Linda Mileshkin, Grant A. McArthur, John Soong, Els M.J.J. Berns, Ross D Hannan, Clare L. Scott, Karen E Sheppard, Richard B Pearson

## Abstract

High-grade serous ovarian cancer (HGSOC) accounts for the majority of ovarian cancer and has a dismal prognosis. PARP inhibitors (PARPi) have revolutionized disease management of patients with homologous recombination (HR) DNA repair-deficient HGSOC. However, acquired resistance to PARPi by complex mechanisms including HR restoration and stabilisation of replication forks is a major challenge in the clinic. Here, we demonstrate CX-5461, an inhibitor of RNA polymerase I transcription of ribosomal RNA genes (rDNA), induces replication stress at rDNA leading to activation of DNA damage response and DNA damage involving MRE11-dependent degradation of replication forks. CX-5461 cooperates with PARPi in exacerbating DNA damage and enhances synthetic lethal interactions of PARPi with HR deficiency in HGSOC-patient-derived xenograft (PDX) *in vivo*. We demonstrate CX-5461 has a different sensitivity spectrum to PARPi and destabilises replication forks irrespective of HR pathway status, overcoming two well-known mechanisms of resistance to PARPi. Importantly, CX-5461 exhibits single agent efficacy in PARPi-resistant HGSOC-PDX. Further, we identify CX-5461-sensitivity gene expression signatures in primary and relapsed HGSOC. Therefore, CX-5461 is a promising therapy alone and in combination therapy with PARPi in HR-deficient HGSOC. CX-5461 is also an exciting treatment option for patients with relapsed HGSOC tumors that have poor clinical outcome.

## Introduction

Ovarian cancer (OVCA) is the major cause of death from gynaecological cancers. The high-grade serous ovarian cancer (HGSOC) subtype accounts for 70-80% of OVCA deaths and overall survival has not changed for several decades (1). HGSOC is characterized by genomic structural variations with relatively few recurrent somatic mutations or driving oncogenes that can be therapeutically targeted. The most frequent genetic alteration occurs in the *TP53* tumor suppressor gene in over 96% of HGSOC cases (1, 2). Further, 50% of HGSOC demonstrate deficiencies in the homologous recombination (HR) DNA repair pathway, most commonly mutations in *BRCA1/2* (3). Aberrations in DNA repair and DNA damage response (DDR) provide a weakness that can be exploited therapeutically (4). As such, tumors with HR deficiency (HRD) exhibit favourable responses to genotoxic chemotherapy and Poly-(ADP-ribose) polymerase (PARP) inhibitors (PARPi) (3–5). Despite significant global research efforts, PARPi are the only class of drugs approved for ovarian cancer in nearly a decade-long drought of new treatments. PARP enzymes are involved in DNA repair through activation of the base excision repair (BER) and alternative end-joining pathways and inhibition of the non-homologous end-joining (NHEJ) pathway (5). More recently PARP was shown to regulate the velocity of DNA replication forks. PARP inhibition increases the speed of fork elongation leading to replication stress, DNA damage and DDR (6). PARP inhibition in cells with HRD is postulated to enhance replication stress and cause accumulation of unrepaired DNA double strand breaks (DSBs), ultimately leading to cell death. Consequently, PARPi are synthetic lethal with HRD (5, 7, 8). PARPi are now utilized as maintenance therapy following complete or partial response to platinum-based chemotherapy in recurrent HGSOC (9, 10). More recently, PARPi has shown substantial benefit with regard to progression-free survival among women with newly diagnosed advanced ovarian cancer with *BRCA1/2* mutations (11). However, resistance to PARPi has been associated with multiple mechanisms including secondary mutations in genes involved in the HR pathway and stabilization of DNA replication forks (12–15). Thus, the development of strategies to overcome resistance to PARPi will provide a significant advancement in the treatment of HGSOC.

Hyperactivation of RNA polymerase I (Pol I) transcription of the 300 copies of ribosomal RNA (rRNA) genes (rDNA) is a consistent feature of cancer cells (16–18). The rDNA repeats are transcribed to produce the 47S pre-rRNA, containing the sequences of 18S, 5.8S and 28S rRNAs, the major ribonucleic acid components of the ribosome. Morphological changes in the nucleoli, the sites of Pol I transcription and ribosome subunit assembly, have been associated with poor cancer prognosis for over 100 years (18). We and others have demonstrated targeting Pol I transcription as a novel approach for cancer treatment (19–22). Inhibition of Pol I transcription using the specific small-molecule inhibitor CX-5461 can selectivity kill cancer cells *in vivo* (19, 20, 23). CX-5461 is currently in phase I clinical trials in patients with haematological malignancies (ACTRN12613001061729) (24) and solid tumors (Canadian Cancer Trials Group, NCT02719977) (25).

We and others have demonstrated that CX-5461 activates p53-independent DDR leading to S-phase delay and G2 cell cycle arrest (25–28). We have shown that by targeting the rRNA gene loci, CX-5461 induces acute Ataxia telangiectasia mutated (ATM) and Ataxia telangiectasia and Rad3 (ATR) kinase signalling in primary fibroblasts prior to the detection of indicators of DNA damage across the genome (27). We also showed that CX-5461 in combination with dual inhibition of checkpoint kinases 1/2 (CHK1/2) downstream of ATM and ATR signalling significantly enhanced the therapeutic outcome of p53-null MYC-driven lymphoma *in vivo* (27). More recently, CX-5461 was shown to exhibit synthetic lethality with *BRCA1/2* deficiency (25). Chronic treatment with CX-5461 in HCT116 colon carcinoma cells was reported to induce stabilization of G-quadruplex DNA (GQ) structures, leading to defects in DNA replication, which presumably require the HR pathway to resolve these defects. However, CX-5461 demonstrated a different spectrum of cytotoxicity compared to the PARPi, olaparib across breast cancer cell lines (25). This suggests that additional mechanisms to HR defects underlie sensitivity to CX-5461.

In this report, we demonstrate that sensitivity to CX-5461 is associated with BRCA mutation and MYC targets gene expression signatures. Further, we demonstrate that CX-5461 exhibits synthetic lethality with HRD in HGSOC cells. While CX-5461 activates ATM/ATR signalling and a G2/M cell cycle checkpoint in HR-proficient HGSOC cells, it induces cell death in HR-deficient HGSOC. Mechanistically, we show that activation of nucleolar ATR is associated with single stranded rDNA coated by replication protein A (RPA) indicating replication stress at rRNA genes and is independent of CX-5461’s ability to stabilize GQ structures. CX-5461 activation of nucleolar ATR leads to activation of DDR and DNA damage involving MRE11-dependent degradation of DNA replication forks. We demonstrate that as single agents CX-5461 and PARPi exhibit different effects on destabilization of replication forks. Importantly, the combination of CX-5461 and PARPi leads to exacerbated DNA damage, pronounced cell cycle arrest and inhibition of clonogenic survival of HR-proficient and HR-deficient HGSOC cells. Remarkably, the growth inhibitory effects of the CX-5461/PARPi combination are independent of HR pathway activity, unveiling a novel CX-5461/PARPi and HRD synthetic lethal axis. Furthermore, the combination of CX-5461 and PARPi leads to significantly improved regression of HR-deficient HGSOC-PDX tumors *in vivo* demonstrating a novel approach to improving the treatment of HR-deficient HGSOC. Importantly, we also provide evidence that CX-5461 has significant therapeutic benefit in PARPi-resistant HGSOC-PDX *in vivo*. Indeed, as CX-5461 activates DNA damage response and destabilises replication forks irrespective of HR pathway status, CX-5461 as a single agent overcomes two well-known mechanisms of resistance to PARPi. Here, we also identify predictive signatures of CX-5461 sensitivity in primary and relapsed ovarian cancer samples highlightling the potential of CX-5461 therapy against primary and aquired chemotherapy- and PARPi-resistant HGSOC.

## Results

### Activity of CX-5461 in OVCA cell lines

The *in vitro* effects of CX-5461 on human OVCA cells were evaluated using a panel of 32 established human OVCA cell lines (29). These cell lines were selected to be representative of a range of histologic OVCA subtypes (Supplementary Table 1). Increasing concentrations (1 nM-10 μM) of CX-5461 were used to assess the concentration of drug that induced a 50% reduction in cell proliferation (GI_50_) at 48 hours (h). The GI_50_ values varied between individual cell lines and ranged from 12 nM for OVCAR3 to 5.17 μM for OV90 (Figure 1A). The cell lines were defined as sensitive to CX-5461 if the GI_50_ was below the geometric median of 363 nM. There was no statistically significant correlation between *TP53* mutation status and sensitivity to CX-5461 (Supplementary Table 1, Figure 1B). The differential GI_50_ effect was not due to the inability of CX-5461 to inhibit Pol I transcription. Both CX-5461-sensitive and - resistant cell lines displayed similar inhibition of Pol I transcription in following 1h treatment with CX-5461, with concentrations that inhibit Pol I transcription by 50% (IC_50_) ranging between 38-285 nM (Figure 1C&D).

**Figure 1.**
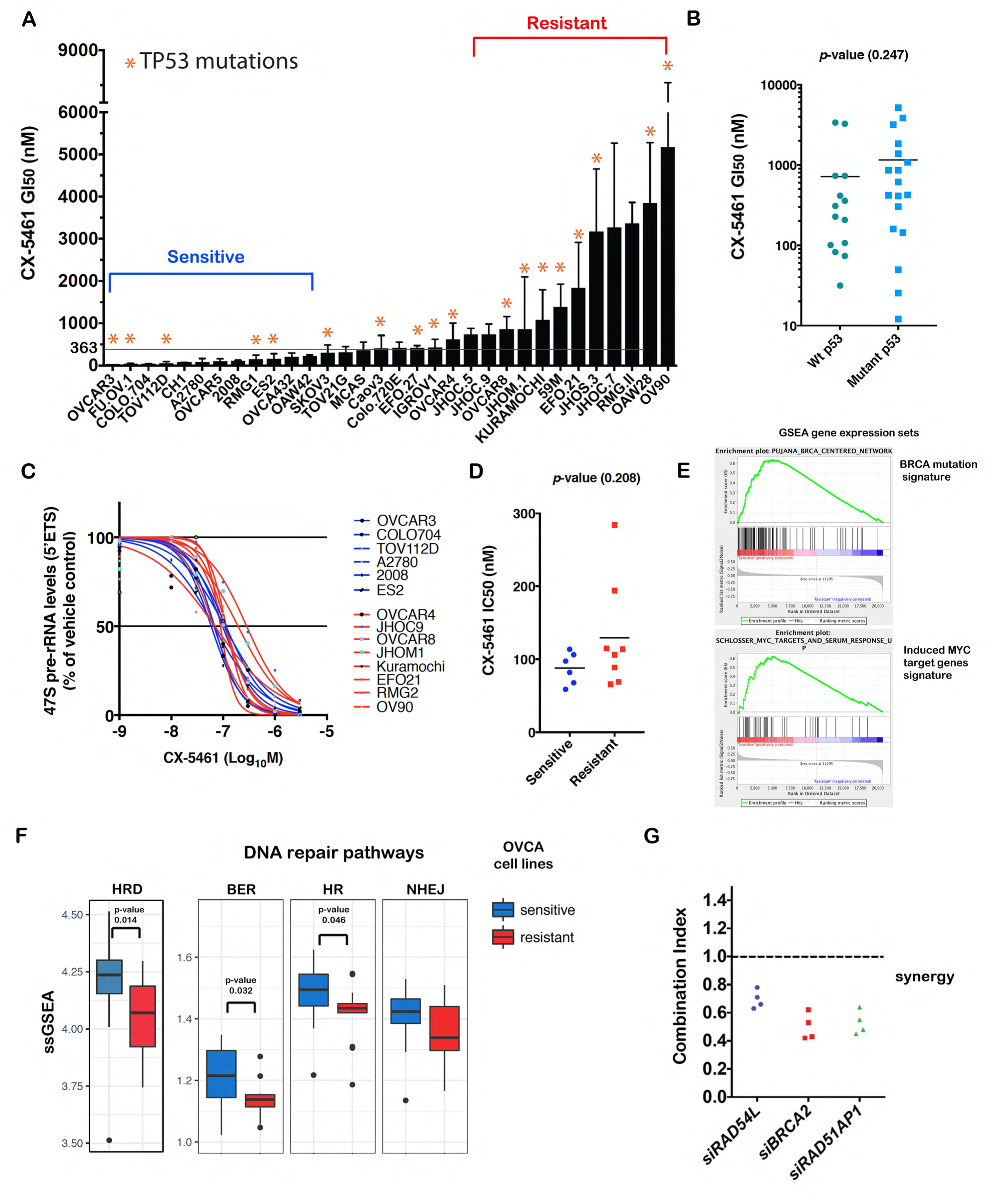
Sensitivity of ovarian cancer cells to CX-5461. **A)** Cell proliferation GI_50_ values (mean ± standard deviation (SD), *n* ≥2) for a panel of ovarian cancer cell lines treated with CX-5461 for 48 hours. * Denote mutated *TP53*. The geometric mean GI50 dose 363 nM is indicated. **B)** CX-5461 exhibit similar activity against p53 wild type and p53 mutant ovarian cancer cell lines. G1_50_ doses of CX-5461 across the OVCA cell line panel as in **A**. Statistical analysis was performed using the Mann-Whitney test. **C)** Dose-response curves of 47S precursor rRNA levels measured by quantitative RT-PCR after 1h treatment with CX-5461 as indicated. Expression levels were normalised to Vimentin mRNA and expressed as fold change relative to vehicle-treated controls. **D)** The inhibitory concentration IC_50_ values (mean ± standard error of the mean (SEM) from **C** are presented, *n* =3. Statistical analysis was performed using the Mann-Whitney test. **E)** GSEA of microarray expression data of 12 CX-5461-sensitive and 11 -resistant cell lines highlighted in **A**. Enrichment plots of the BRCA network and MYC targets gene sets identified to be enriched in the CX-5461-sensitive OVCA cell lines are shown. **F)** ssGSEA was utilized to obtain the level of activity of pathways in individual samples. Genes in each sample were ranked according to their expression levels, and a score for each pathway was generated based on the empirical cumulative distribution function, reflecting how highly or lowly genes of a pathway were found in the ranked list. Statistical significance of the ssGSEA scores was obtained using two-sided Wilcoxon tests. Benjamini-Hochberg correction was subsequently used to account for multiple testing. **G)** The combination of CX-5461 and siRNAs targeting HR genes in OVCAR4 cells synergistically inhibits proliferation. The graphs are bliss plots of HR pathway components where each dot represents an individual siRNA duplex. Cell proliferation was measured by cell count using DAPI staining and imaging using Cellomics. A combination index of CI < 1 indicates synergy, CI > 1 indicates antagonism and CI = 1 indicates additive effect. Shown are four independent siRNAs per gene.

### CX-5461 exhibits synthetic lethality with HR deficiency in HGSOC

To interrogate the molecular mechanisms underlying CX-5461 sensitivity in ovarian cancer, gene expression profiles for 12 CX-5461-sensitive and 11 -resistant cell lines were generated (Figure 1A). Gene set enrichment analysis (GSEA) identified BRCA mutation and MYC targets gene expression signatures to correlate with sensitivity to CX-5461 *in vitro*. (Figure 1E, Supplementary Figure 1). Indeed, we observed a significant enrichment of HRD gene signature (30) in the CX-5461-sensitive cell lines (Figure 1F) suggesting functional defects in the HR pathway underlying sensitivity to CX-5461. Defects in the HR pathway and/or replicative stress can result in transcriptional rewiring of DNA repair pathways (31). We therefore examined the expression levels of genes involved in BER, HR, and NHEJ DNA repair pathways by performing single sample GSEA (ssGSEA) (32, 33) (Figure 1F), which determines the activity level of pathways by measuring the consistency in the expression levels of its individual genes. Our analysis identified the expression of genes involved in HR and BER, but not NHEJ, to be significantly upregulated in CX-5461-sensitive compared to CX-5461-resistant OVCA cell lines. This is consistent with other studies demonstrating HR-related genes to be highly expressed in cancer cells that harbor low HR efficiency (31). As such, the data suggest synthetic lethal interaction between defects in HR and/or BER with CX-5461. Further, CX-5461 and PARPi showed little overlap in spectrum of sensitivity across HGSOC cell lines (Supplementary Figure 2), suggesting that additional mechanisms to HRD confer sensitivity to CX-5461. As sensitivity to CX-5461 was also enriched with a MYC targets gene expression signature (Figure 1E), our data suggest CX-5461 may provide therapeutic benefit in the high-MYCN HGSOC subtype classified with elevated functional MYC activity and poor progression free survival (34, 35).

To confirm CX-5461 synthetic lethal interaction with HRD, we tested the effects of short interfering RNAs (siRNAs) targeting HR genes on the efficacy of CX-5461 and demonstrated strong synergistic inhibition in HGSOC OVCAR4 cells, reported to closely resemble the genomic profile of HGSOC tumors (36) (Figure 1G). Further, we tested CX-5461’s interaction with HRD in isogenically matched HR-proficient OVCAR8 cells and a derivative HR-deficient RAD51C knockout (KO) OVCAR8 cell line as well as a cell line derived from a chemo-naïve HGSOC-PDX (#62, WEHICS62) with reduced capacity to form RAD51 foci in response to ionizing radiation (IR) damage due to *BRCA1* promoter hypermethylation (37) (Supplementary Figure 3A&B). Both HR-deficient cell lines exhibited increased sensitivity to growth inhibition by CX-5461 at the low doses of 10 nM and 100 nM compared to OVCAR8 cells (Figure 2A). In HR-proficient OVCAR8 cells, CX-5461-mediated inhibition of proliferation was associated with a pronounced G2/M cell cycle arrest, cytokinesis failure and multinucleation (8N DNA content) suggesting that CX-5461 induces defects in DNA replication leading to mitotic DNA damage (Figure 2B&C). In comparison, HR-deficient RAD51C KO OVCAR8 cells underwent cell death at 100 nM and 1 μM CX-5461 further confirming synthetic lethal CX-5461/HRD interaction (Figure 2B&C).

**Figure 2.**
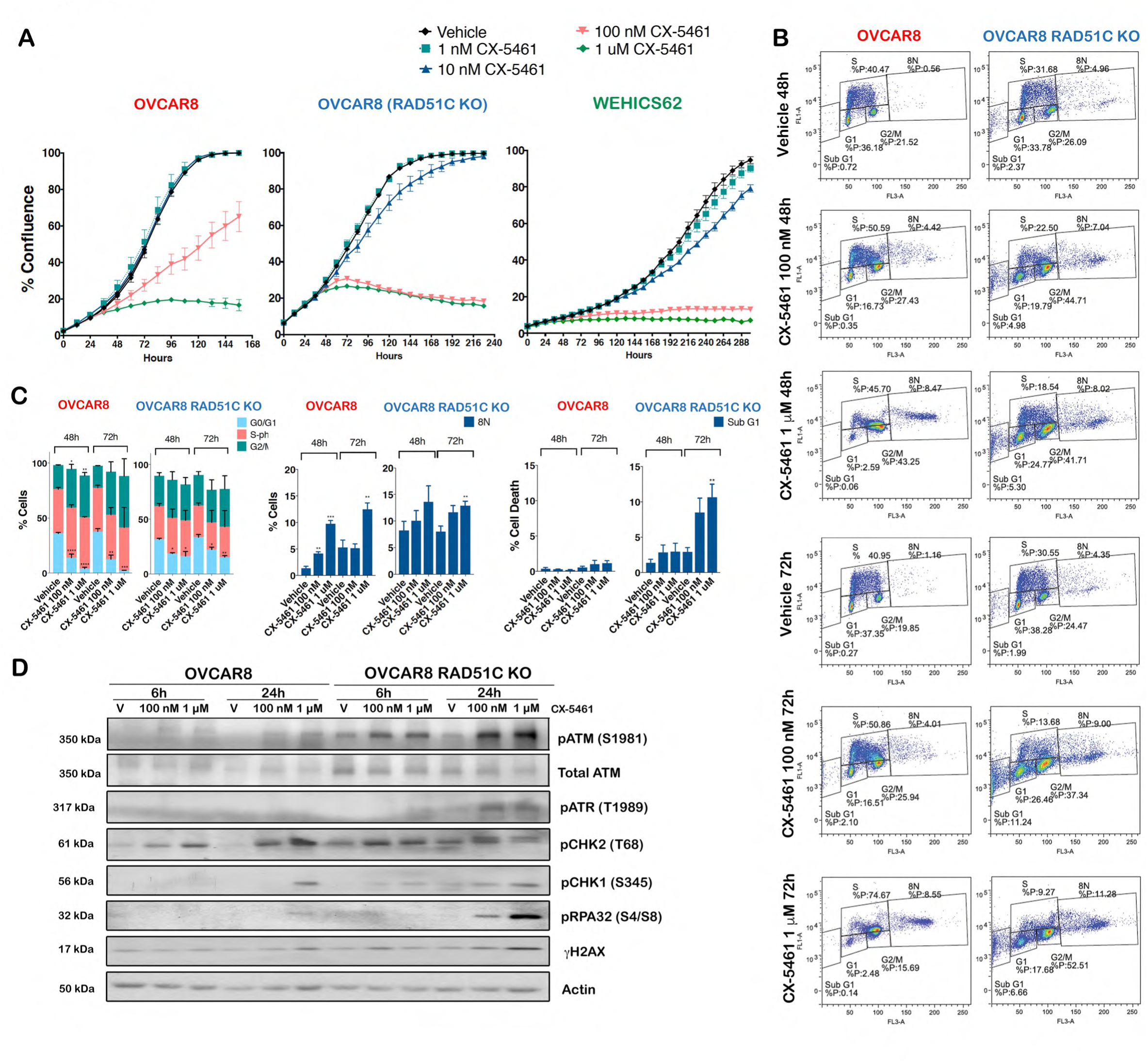
CX-5461 is synthetic lethal with HRD in HGSOC. **A)** *In vitro* CX-5461 dose response proliferation time course, assessed by cell confluency using IncuCyte ZOOM of OVCAR8, OVCAR8 derivative with RAD51C KO and the WEHICS62 HGSOC cell lines. Representative of *n* ≥ 2, mean ±SEM of 5 technical replicates. **B)** Cell cycle analysis of cells treated with vehicle 100 nM or 1μM CX-5461 for 48h and 72h and labelled with BrdU for 30 min prior to harvest. Analytical FACS analysis showing BrdU incorporation as a function of DNA content. The boxes represent S-phase BrdU-labelled population G0/G1 and G2M, Sub G0/G1 cell populations and cells with > 4n content. **C)** Histogram plots for total cells stained from the corresponding cell populations shown in **B**, (*n* = 3), error bars represent mean ± SEM. Statistical analysis was performed using one-way ANOVA multiple comparisons, \**p* < 0.05, \*\**p*-value < 0.01, \*\*\**p*-value < 0.001 relative to corresponding vehicle treated control. **D)** Western blot analysis of cells treated with either vehicle, 100nM or 1μM CX-5461 for 6h and 24h. Representative of *n* =3.

We have previously shown that CX-5461 activates p53-independent ATM/ATR-mediated S-phase and G2/M cell cycle checkpoints (27). We therefore investigated CX-5461 mediated activation of ATM/ATR signalling in HGSOC cell lines (Figure 2D). In agreement with our previous findings (27), 1 μM CX-5461 induced ATM/ATR signalling, as indicated by increased phosphorylation of CHK2 on T68 and CHK1 on S345 in HR-proficient OVCAR8 cells. However, enhanced ATM and ATR activation following treatment with 100 nM and 1 μM CX-5461 were observed in HR-deficient RAD51C KO cells as well as increases in S4/S8 phosphorylation of RPA32, an immediate responder to chromatin defects that protects single-stranded DNA (ssDNA). RPA32 S4/8 phosphorylation is mainly catalysed by ATM and DNA-dependent protein kinase (DNA-PK) at persistently stalled replication forks (38). Furthermore, CX-5461 led to higher levels of γH2AX, a marker of DNA damage in HR-deficient cells compared to HR-proficient cells. Thus, CX-5461 induces replication stress in HR-proficient cells associated with increased percentage of multinucleated cells while HR-deficient cells undergo cell death following CX-5461 due to exacerbated replication stress and DNA damage. Altogether, these findings demonstrate that CX-5461 exhibits strong antiproliferative effects in OVCA cells with HR defects sensitizing HGSOC cells to CX-5461-mediated cell death, consistent with the BRCA-mutated and HRD gene expression signature predicting OVCA cells’ sensitivity to CX-5461 (Figure 1E&F). However, CX-5461-sensitivity gene expression signatures suggest additional mechanisms to HRD such as defects in BER and increased MYC activity may also confer sensitivity to CX-5461 (Figure 1E&F).

### CX-5461 induces nucleolar-specific DDR

Prolonged CX-5461 treatment in HCT116 colon carcinoma cells was reported to be associated with stabilization of G-quadruplex DNA (GQ) structures leading to defects in DNA replication and activation of DDR (25). These non-canonical DNA structures can influence DNA replication, transcription, and protein sequestration and are abundant at the rRNA genes due to their repetitive and highly transcribed nature. To examine the mechanism of activation of ATM/ATR signalling, we investigated the effects of acute CX-5461 treatment on the formation of GQ structures in OVCAR8 cells and RAD51C KO derivative cells (Figure 3A&B) as well as two additional HGSOC cell lines, OV90 and OVCAR4 (Supplementary Figure 4A&B). CX-5461 did not affect GQ stabilization in HR-proficient or HR-deficient OVCAR8 cells nor OVCAR4 cells compared to a *bona fide* GQ stabilizer (TMPyP4) (Figure 3 A&B). In contrast, 1h CX-5461 treatment of OV90 cells induced GQ stabilization with similar kinetics to TMPyP4 (Supplementary Figure 4A&B), indicating context-dependent effects of CX-5461 on GQ stabilization. However, at 1h and 3h post-treatment, TMPyP4 did not activate DDR while CX-5461 induced DDR signalling in HGSOC cell lines (Supplementary Figure 4C-E). While we did not investigate CX-5461’s effect on GQ stabilisation after 24h treatment as reported in (25), our data clearly demonstrate that CX-5461-mediated activation of DDR is independent of its role in GQ stabilization.

**Figure 3.**
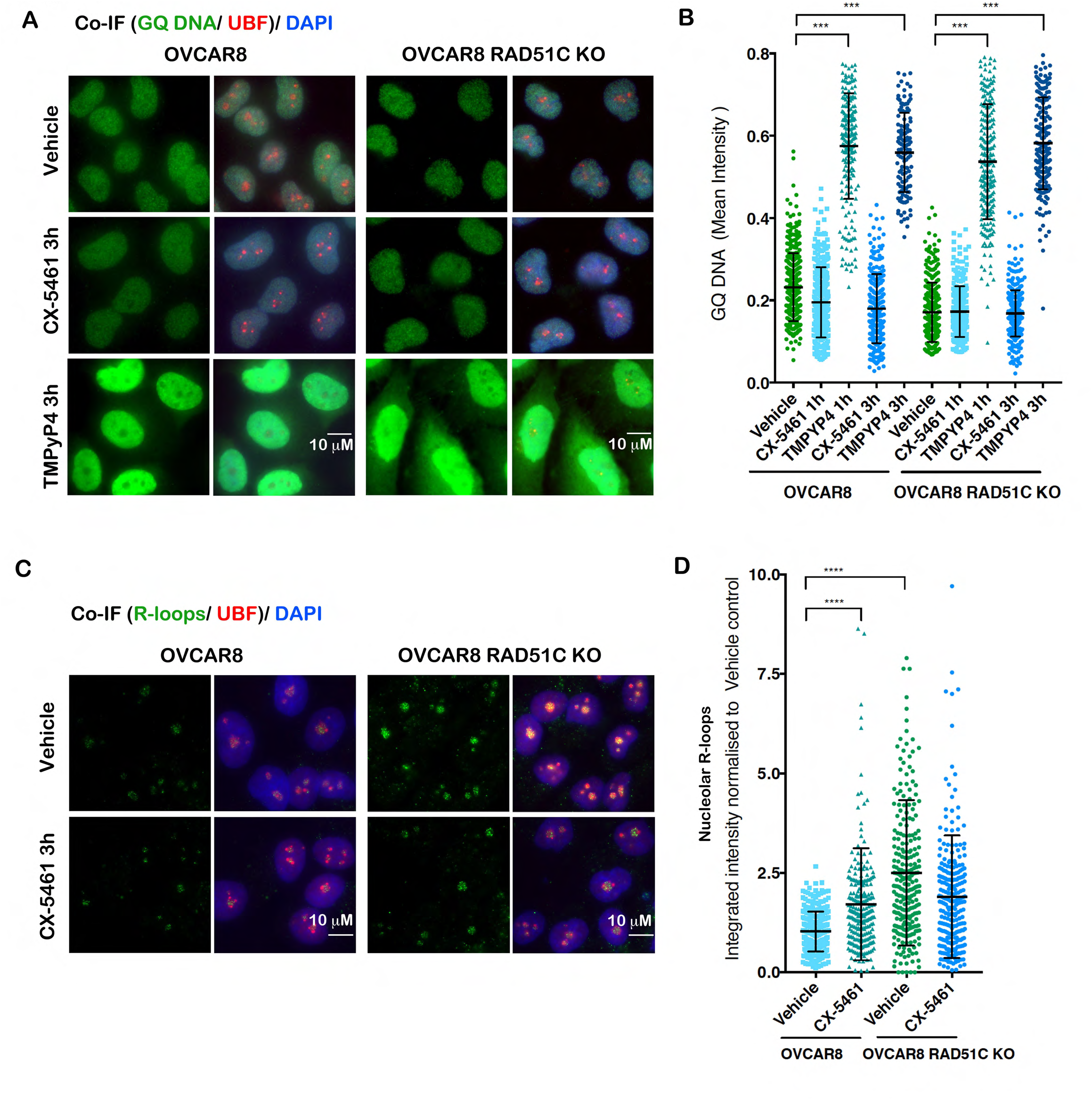
CX-5461 induces R-loops stabilization in HR-proficient OVCAR8 cells. **A)** Co-immunofluorescence (Co-IF) of GQ DNA and UBF as a nucleolar marker in OVCAR8 cells and RAD51C KO OVCAR8 cells treated with either vehicle, 1 μM CX-5461 or 10 μM TMPyP4 for 3h. Representative images of *n* =3. **B)** Quantitation of GQ DNA immunofluorescence. Signal intensities were analyzed using Cell Profiler. Error bars represent mean ± SD, *n* =3, >200 cells per condition. Statistical analysis of the increase in GQ signal intensity was performed using Kruskal-wallis test, \*\**p*-value < 0.01, \*\*\**p*-value < 0.001 compared to vehicle control. **C)** Co-IF analysis of R-loops and UBF in cells treated with vehicle or 1 μM CX-5461 for 3h. **D)** Quantitation of R-loops signal intensity was performed using Cell Profiler *n* =2, >200 cells per condition. Statistical analysis was performed using Kruskal-wallis test, \*\*\**p*-value < 0.001, \*\*\*\**p*-value < 0.0001.

We next examined the formation of RNA:DNA hybrids (R-loops), which are by- products of Pol I transcription. Stabilization of these three-stranded structures of nucleic acids consisting of a DNA-RNA hybrid and displaced single-stranded DNA (ssDNA) are known to obstruct DNA replication and activate DDR (39). As Pol I transcription initiation, elongation and rRNA processing are tightly co-regulated, we therefore investigated the effects of CX-5461-mediated inhibition of Pol I transcription initiation on nucleolar R-loops formation. To mark nucleolar R-loops we performed co-IF for the upstream binding transcription factor (UBF), which localizes to decondensed rDNA chromatin (40, 41). A striking change in nucleolar R-loop staining was observed, from being weak and diffuse in control OVCAR8 and OVCAR4 cells to a focal pattern with more intense foci in CX-5461-treated cells (Figure 3C&D and Supplementary Figure 5A-C). This suggests that CX-5461-induced Pol I displacement from the promoters of rRNA genes (27, 42) is associated with the formation of stable rRNA transcript:single-strand rDNA hybrids. Intriguingly, RAD51C KO OVCAR8 cells exhibited high level of stable R-loops compared to parental cells (Figure 3 C&D) suggesting transcription-replication conflict stress at rRNA genes. This indicates loss of HR activity contributes to R-loops formation and these structures remain at high levels post-treatment with CX-5461 suggesting persistent defects in nucleolar chromatin may contribute to the CX-5461 and HRD synthetic lethal interaction.

To further analyse the mechanisms of activated nucleolar DDR, we examined whether CX-5461 induces ATR at the nucleoli (Figure 4A). Indeed, phosphorylated ATR T1989 was detected following CX-5461 treatment at the periphery of nucleolar R-loops, co-localizing with UBF in CX-5461 treated HR-proficient OVCAR8 and OVCAR4 cells (Figure 4A&B, Supplementary Figure 5D-F & Supplementary Figure 6A). Localization of ATR to the periphery of the nucleoli is in agreement with damaged rDNA movement to anchoring points at the nucleolar periphery where it is more accessible to DNA repair factors (43). However, activation of nucleolar ATR in response to CX-5461 was not restricted to S-phase (EdU positive) cells and was significantly induced in HR-defective cells (Figure 4A&B). These data suggest that ATR is activated in response to CX-5461-induced defects at rDNA chromatin, as opposed to DNA replication defects and that these rDNA chromatin defects are significantly exacerbated in the absence of intact HR.

**Figure 4.**
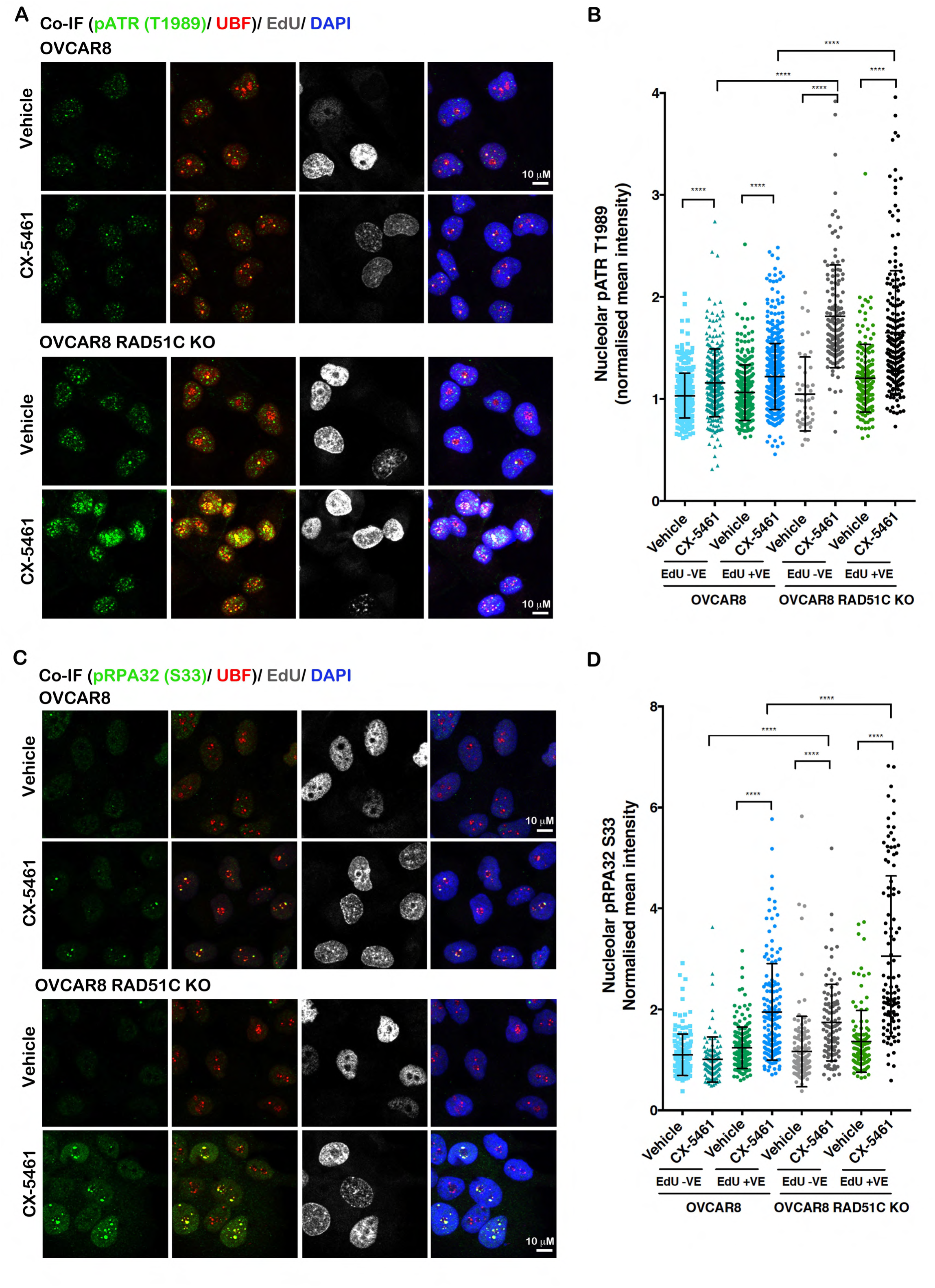
CX-5461 activates nucleolar-specific DDR. **A)** Co-IF analysis of pATR(T1989) and UBF in cells labelled with EdU and treated with vehicle or 1 μM CX-5461 for 3h. **B)** Quantitation of signal intensity of the colocalized regions was performed using Cell Profiler and normalized to the median of vehicle treated controls. Statistical analysis was performed using Kruskal-wallis test, *n* =3, >165 cells per condition, \*\*\*\**p*-value < 0.0001. **C)** Co-IF analysis of pRPA32 (S33) and UBF in cells labelled with EdU and treated with vehicle or 1 μM CX-5461 for 3h. **D)** Quantitation of signal intensity of the colocalized regions was performed using Cell Profiler and normalized to the median of vehicle treated controls. Statistical analysis was performed using Kruskal-wallis test, *n* =2, >290 cells per condition, \*\*\*\**p*-value < 0.0001.

RPA-coated ssDNA recruits and activates numerous DNA repair and cell cycle checkpoint regulators including ATR. ATR phosphorylates RPA32 at S33 immediately after fork stalling (38). We therefore investigated phosphorylation of RPA32 at S33 upon CX-5461 treatment and observed significant localization of S33 phosphorylated RPA32 at the nucleoli in both HR-proficient and HR-deficient cells (Figure 4C-D). While CX-5461-mediated recruitment of nucleolar RPA was specific to the S-phase population in HR-proficient cells, nucleolar RPA recruitment was independent of the cell cycle stage in HR-deficient cells. This is consistent with HR-mediated repair of rRNA genes occurring throughout the cell cycle (43). Thus, our data implicates HRD in exacerbating CX-5461-mediated replication stress at rRNA genes and this may underpin CX-5461’s synthetic lethal interaction with HRD.

### CX-5461 induces MRE-11-mediated replication stress and cooperates with PARPi in exacerbating replication stress

Our data demonstrated a robust increase in pRPA32 S4/S8 and γH2AX levels by western blotting in HR-deficient cells following 24h treatment with CX-5461 compared to the HR-proficient counterparts (Figure 2D). We next examined γH2AX foci formation after 3h treatment with CX-5461 at which time ATR and RPA were recruited to the nucleoli. CX-5461 induced global γH2AX foci only in the EdU-positive population of OVCAR8 cells, suggesting that, by contrast to ATR activation, CX-5461-induced DNA damage is associated with DNA replication (Figure 5A&B). Thus, CX-5461 induces replication-dependent DNA damage in HR-proficient cells. Strikingly, HR-deficient RAD51C KO OVCAR8 cells exhibited significantly high levels of γH2AX foci in the EdU-positive and -negative populations compared to HR-proficient cells (Figure 5A&B), implicating cooperation between loss of RAD51 function with CX-5461-mediated replication stress in exacerbating DNA damage. Furthermore, we observed a significant increase in the percentage of HR-deficient cells exhibiting γH2AX foci after 24h treatment with CX-5461 compared to HR-proficient cells (Figure 5C), indicating that the HR pathway is required for repairing CX-5461-mediated replication stress.

**Figure 5.**
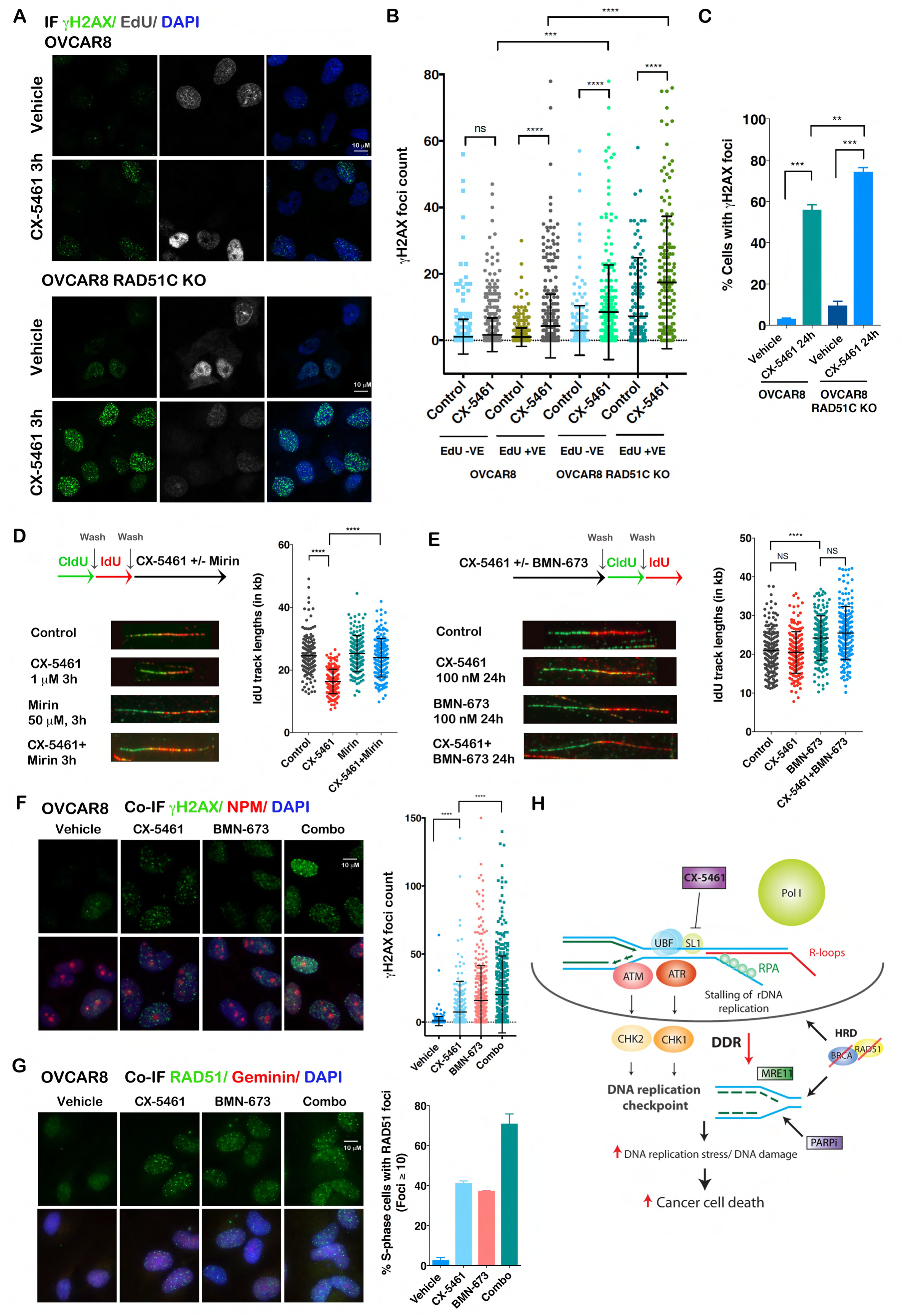
CX-5461-induces replication-dependent DNA damage in HR-proficient HGSOC cells. **A)** Co-IF analysis of γH2AX in cells labelled with EdU and treated with vehicle or 1 μM CX-5461 for 3h. **B)** Quantitation of foci count was performed using Cell Profiler. Statistical analysis was performed using Kruskal-wallis test, *n* =3, >180 cells per condition, ***< 0.001, \*\*\*\**p*-value < 0.0001. ns denotes not significant p-value. **C)** Quantitation of the percentage of OVCAR8 and OVCAR8 RAD51C KO cells with > 10 γH2AX foci. Cells were treated with vehicle or 100 nM CX-5461 for 24h, mean ± SEM, *n* =3 for OVCAR8 and *n* =2 for OVCAR8 RAD51C KO cells. Statistical analysis was performed using one-way ANOVA multiple comparisons, \**p* < 0.05, \*\**p*-value < 0.01, \*\*\**p*-value < 0.001. **D)** IdU track length is reduced by CX-5461 through MRE11-dependent mechanism. Schematic of CIdU and IdU pulse-labelling (top). OVCAR8 cells were sequentially labelled and either processed or treated with 1 μM CX-5461, 50 mM mirin or CX-5461+ mirin for 3h. Fibres were processed for DNA fibre analysis; *n*=2. Replication Fork length was calculated based on the length of the IdU tracks measured using Image J software. At least 150 replication tracks were analysed per experiment. IdU track lengths (in um) were converted to kb (1kb=2.59 um). p-value was calculated by Kruskal-Wallis test, \*\*\*\**p*-value < 0.0001. **E)** DNA fibre analysis of OVCAR8 cells pre-treated with 100 nM CX-5461, 100 nM BMN-673 or in combination, washed then sequentially labelled with CldU and IdU as indicated in the schematic (top). Fibres were processed an analyzed as described above. At least 150 replication tracks were analysed per experiment, *n*=2. p-value was calculated by Kruskal-Wallis test, \*\*\*\**p*-value < 0.0001. ns denotes no significant change (p-value greater than 0.05). **F)** CX-5461 and BMN-673 induce markers of DNA damage in OVCAR8 cells. OVCAR8 cells were treated with vehicle, 100 nM CX-5461, 100 nM BMN-673 or the combination for 24h. Co-IF for γH2AX and nucleophosmin (NPM), as a nucleolar marker was performed and representative images of *n*=3 are shown. Quantitation of foci count was performed using Cell Profiler. Statistical analysis was performed using Kruskal-wallis test, *n* =3, >170 cells per condition, ****p-value < 0.0001. **G)** OVCAR8 cells we treated as in (F). Co-IF of RAD51 and Gemini was performed. Representative images and quantitation of the percentage of Gemini +ve cells with > 10 RAD51 foci is shown. Graphs represent mean ± SD, *n* =2. **H)** A schematic of molecular response to CX-5461. CX-5461 inhibits the Pol I transcription complex by binding the selectivity complex 1 (SL-1) and preventing Pol I from binding to the rRNA gene promoters. Displacement of Pol I and inhibition of Pol I transcription initiation are associated with R-loops formation, recruitment of RPA to single strand rDNA, replication stress and activation of DDR at the nucleoli. The net result of active nucleolar DDR is the induction of S-phase and G2/M cell cycle arrest, which is associated with a halt in DNA replication, termed the DNA replication checkpoint. Activation of DDR involves MRE11-dependent degradation of replication forks. The HR pathway and PARP activity are necessary to counteract DNA replication stress. Thus, CX-5461 co-operates with HRD and inhibition of PARP activity in exacerbating replication stress and DNA damage promoting cell death.

BRCA1/2 and RAD51 play major roles in replication fork stabilization following replication stress by preventing nucleolytic degradation of replication forks by the nuclease MRE11 (44). Stabilization of replication forks is one of the mechanism associated with resistance to PARPi in BRCA-deficient HGSOC. We therefore performed DNA fibre analysis to investigate the effect of CX-5461 on fork stabilisation (Figure 5D & Supplementary Figure 6B) in OVCAR8 cells. Nascent replication tracks were sequentially labelled with CldU and IdU before treatment with CX-5461 for 3h. The data shows CX-5461 treatment causes an overall decrease in track length, suggesting degradation of replication forks upon induction of DDR by CX-5461. This was rescued by co-treatment with the MRE11 inhibitor mirin, confirming inhibition of the MRE11 nuclease can rescue CX-5461-mediated fork destabilisation. We next assessed whether DNA damage induced by CX-5461 treatment affects fork progression by pre-treating cells with CX-5461 for 24h and then pulse labelled with both analogs (Figure 5E). Pre-treatment with CX-5461 had no effect on fork length suggesting CX-5461 does not cause any lesions that could prevent fork restarting or progression. On the other hand, the PARPi talazoparib (BMN-673) increased fork progression in agreement with a recent report implicating PARPi mediated acceleration fork elongation as a novel mechanism for replication stress and DNA damage (6). Thus, our data demonstrate that CX-5461 and PARPi cause replication stress via different effects on fork destabilisation indicating independent synthetic lethal interactions with HRD. Indeed, the combination of CX-5461 and BMN-673 led to a significant increase in γH2AX foci formation (Figure 5F) suggesting their cooperation in exacerbating replication stress and DNA damage. As the HR pathway is required to counteract replication stress by stabilizing stalled replication forks, we investigated the possibility of CX-5461 destabilising replication forks via inhibiting HR. We found CX-5461 induced RAD51 foci formation, which indicated loading of RAD51 onto DSBs while the combination of CX-5461 and BMN-673 led to further enhanced RAD51 foci formation (Figure 5G). Thus, CX-5461-mediated DDR activates HR, up to the stage of RAD51 loading and its cooperation with BMN-673 in exacerbating replication stress is independent of the HR pathway. Taken together, our data are consistent with a model (Figure 5H) by which CX-5461 inhibits Pol I recruitment leading to rDNA chromatin defects including stabilization of R-loops, ssDNA and replication stress at the rRNA gene loci. The net effect of CX-5461-induced targeted DDR is destabilization of replication forks via MRE11 activity leading to global replication stress and DNA damage. CX-5461 and PARPi exhibit different effects on fork destabilisation and in combination they exacerbate replication stress leading to accumulation of DNA damage and subsequent cell death. HRD potentiates CX-5461-mediated nucleolar and global DDR and this underpins the PARPi/ CX-5461/ HRD synthetic lethal interactions.

### CX-5461 cooperates with PARPi in inhibiting HGSOC cell growth and survival

Since CX-5461 in combination with BMN-673 lead to increased replication stress and the CX-5461-sensitivity spectrum in HGSOC cell lines differs to PARPi (Supplementary Figure 2), we hypothesized that combining CX-5461 with PARPi or other DNA repair and DDR inhibitors (DDR therapy) may improve the efficacy of treating HGSOC. We therefore performed a focused/boutique drug screen in the HR-proficient OVCAR4 cells for DNA repair and DDR inhibitors that may cooperate with CX-5461 to enhance growth arrest (Figure 6A & Supplementary Figure 3C). Inhibitors of ATM (ATMi: KU55933), ATR (ATRi: VE-821), PARP (BMN-673), the platinum-based chemotherapy drug cisplatin, the mTORC1 inhibitor everolimus and the selective inhibitor of BCL-2 ABT-199 all demonstrated growth inhibitory effects as single agents. However, the combination of GI_20_ dose of CX-5461 with BMN-673 and VE-821 (ATRi) showed the most enhanced proliferative arrest compared to single-agent effects and compared to combinations with other compounds (Figure 6A). ATR inhibition leads to degradation of stalled replication forks. Thus, CX-5461 may preferentially cooperate with PARPi and ATRi in destabilising replication forks and enhancing replication stress. We next examined the cooperation between CX-5461 and BMN-673 or ATRi in inducing cell death in 3 additional HR-proficient HGSOC cell lines (Supplementary Figure 3C) and identified robust and significant interactions between BMN-673 and CX-5461 in inducing cell death compared to the ATR inhibitor (VE-821) (Figure 6B). In addition, dose-response curves of BMN-673 in the presence or absence of CX-5461 confirmed CX-5461’s strong interaction with BMN-673 (Figure 6C). Further, the combination of CX-5461 and BMN-673 led to enhanced G2/M cell cycle arrest, enhanced inhibition of cell proliferation and significantly reduced clonogenic survival of HR-proficient OVCAR8 and HR-deficient RAD51C KO OVCAR8 cells and OV90, the most CX-5461-resistant HGSOC cell line (Figure 6E&F).

**Figure 6.**
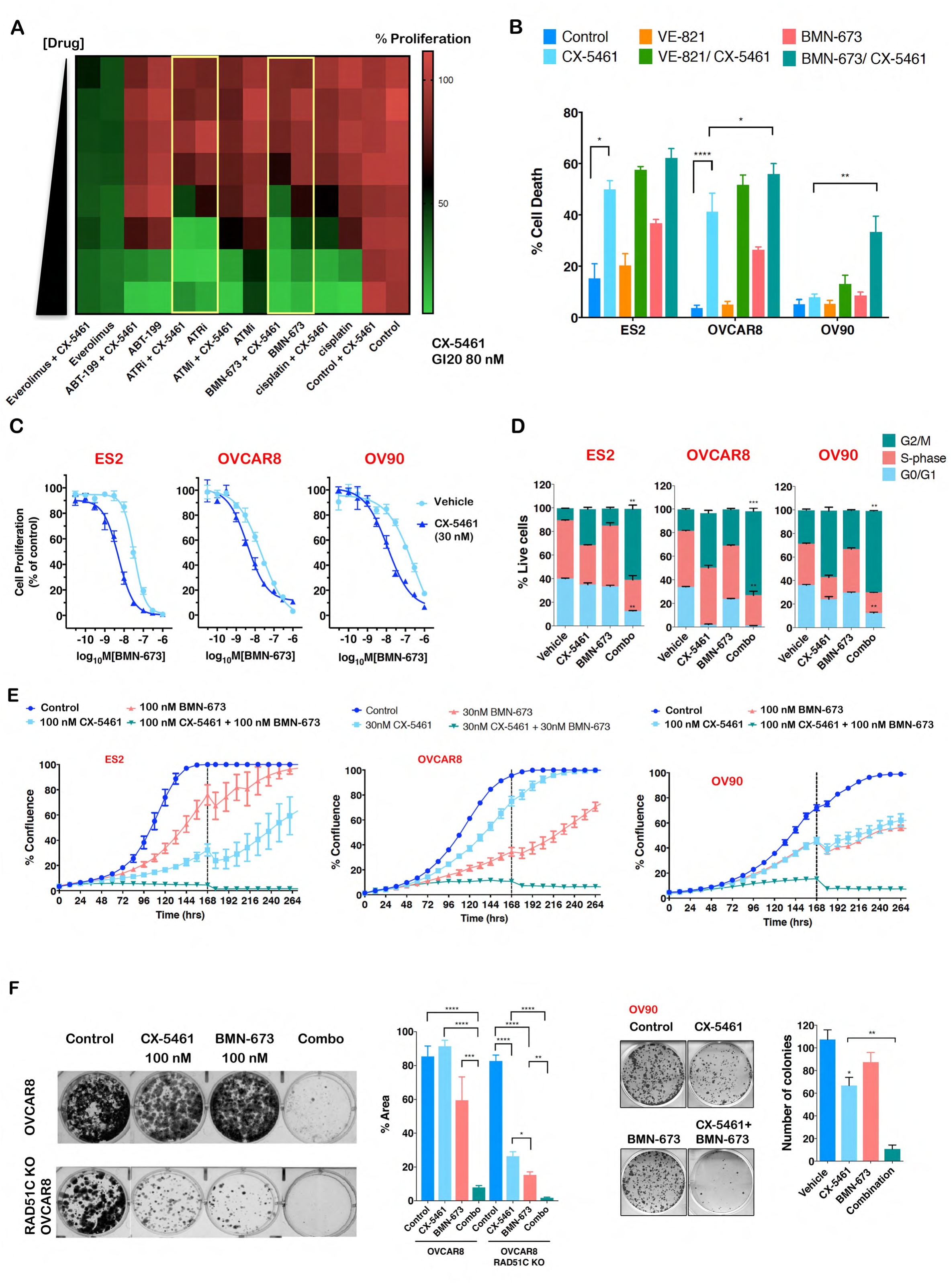
CX-5461 cooperates with PARPi in inhibiting HGSOC cell growth. **A)** Mini drug screen performed in HR-proficient OVCAR4. Cells were treated with increasing doses of cisplatin (0-1.11 μM), PARPi (BMN-673, 0-0.11 μM), ATMi (KU55933, 0– 1.11 μM), ATRi (VE-821, 0–1.11 μM), ABT-119 (0–1.11 μM) or Everolimus (0-0.11 μM) in the absence or presence of 80 nM (GI_20_) CX-5461. Color-coding denotes the level of proliferation as measured by DAPI staining and imaging using Cellomics (green denotes reduced proliferation). Dose response of single drug treatments were corrected for vehicle treatment control and the combination was corrected for response in the presence of 80 nM CX-5461, the average values of *n*=5 are presented. The combination of CX-5461 with BMN-673 or ATRi showed enhanced proliferation arrest, highlighted by yellow boxes. **B)** Quantitation of cell death, determined by PI staining and SubG1 DNA content analysis of cell treated with vehicle, 100 nM CX-5461, 1 μM VE-821, 100 nM BMN-673 or in combination for 7 days (mean ± SEM), *n* =3. Statistical analysis was performed using two-way ANOVA multiple comparisons, \**p* < 0.05, \*\**p*-value < 0.01. **C)** Representative BMN-673 dose response curves in the absence (light blue) or presence (dark blue) of 30 nM CX-5461. Cell proliferation was measured using SRB assays at 5 days post-treatment. Dose response curves for BMN-673 only, were corrected for DMSO treatment control and the combination was corrected for response in the presence of 30 nM CX-5461. Representative of *n* =3. **D)** Cell cycle analysis of cells treated with vehicle, 1μM CX-5461, 100 nM BMN-673 or in combination for 72h and labelled with BrdU for 30 min prior to harvest. The percentage of live BrdU-labelled S-phase and BrdU-negative G1/G0 and G2/M populations was determined using Flowlogic software. *n* =3, mean ± SEM, statistical analysis was performed using two-way ANOVA multiple comparisons, \**p* < 0.05, \*\**p*-value < 0.01, \*\*\**p*-value < 0.001 relative to corresponding CX-5461-treated controls. **E)** *In vitro* proliferation time course assessed by cell confluency using IncuCyte ZOOM of ES2, OVCAR8 and OV90 cells following treatment with CX-5461, BMN-673 alone and in combination as indicated. Representative of *n* ≥ 3, mean ± SEM of 5 technical replicates. Dashed lines denote resupplement of media with drugs. **F)** CX-5461 and BMN-673 cooperate in inhibiting clonogenic survival of OVCAR8, RAD51C KO OVCAR8 and OV90 cells. Representative of *n*=3, mean ± SEM. Statistical analysis was performed using one-way ANOVA multiple comparisons, \**p*-value < 0.05, \*\**p*-value < 0.01, \*\*\**p*-value < 0.001, \*\*\*\**p*-value < 0.0001.

Next, we investigated the effects of combining CX-5461 with BMN-673 on Pol I transcription rates based on the fact that nucleolar PARP1 is implicated in regulation of rDNA heterochromatin (45). Reverse transcription real-time PCR was used to measure 47S rRNA precursor levels following treatment of OVCA cells with single agent CX-5461 or BMN-673 or their combination (Supplementary Figure 7). After 3h treatment with CX-5461, we detected significant decreases in rRNA precursor levels using primers specific to the external spliced region (5’ETS) (+413-521bp) relative to the transcription start site (TSS). However, the combination of CX-5461 with BMN-673 did not further decrease rRNA abundance compared to CX-5461-treated samples. Therefore, our data suggest CX-5461 cooperates with PARPi in inducing cell death and inhibiting survival of HR-proficient and HR-deficient HGSOC cells by exacerbating replication stress and DNA damage in HGSOC (Figure 5H) as opposed to enhancing inhibition of Pol I transcription.

### CX-5461 has significant therapeutic efficacy against HR-deficient olaparib-sensitive and olaparib-resistant HGSOC *in vivo*

We next investigated the potential of CX-5461 and PARPi interaction *in vivo* in a *BRCA2*-mutated, HR-deficient post one line of platinum treatment HGSOC-PDX (#19B). The administration of CX-5461 and olaparib as single agents resulted in stable disease and a statistically significant survival benefit (median time to harvest (TTH) for CX-5461 treatment 53 days, olaparib 67 days vs vehicle 22 days, *p*-values 0.00285 and 0.00285 compared to vehicle, respectively) (Figure 7A). Remarkably, co-treatment of CX-5461 and olaparib was well-tolerated (Supplementary Figure 7B) and resulted in dramatic durable regression with reductions in tumor volumes indicating partial remission (defined as reduction in tumor volume of > 30% from baseline) with survival lasting more than 100 days (median TTH 113 days, *p*-values 0.00692 compared to CX-5461 single agent treatment). Thus, CX-5461 has additional activity to olaparib and the strong cooperation between CX-5461 and PARPi provides a major improvement for overall survival in HR-deficient HGSOC. This is in agreement with CX-5461 and PARPi having different mechanisms of response and spectrum of sensitivity in HGSOC cell lines (Supplementary Figure 2).

**Figure 7.**
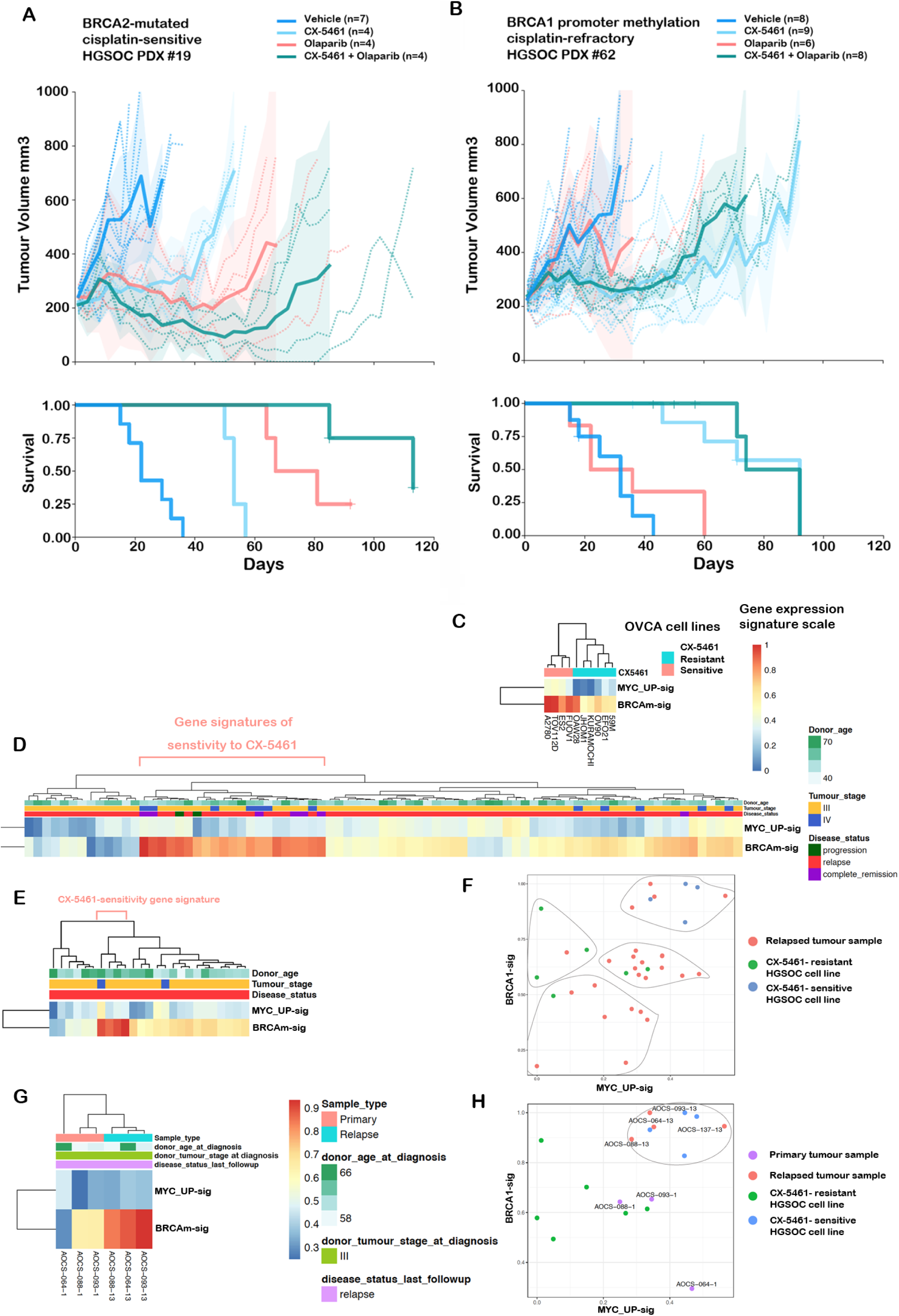
**A)** Responses observed in post-platinum treated BRCA2-mutant PDX#19 HGSOC patient-derived xenografts (PDX) **(A)** and PDX #62 with *BRCA1* promoter methylation **(B)** to CX-5461 and olaparib treatment *in vivo*. Recipient mice bearing the PDX were randomized to treatment with vehicle, 40 mg/kg CX-5461 twice a week, 50 mg/kg olaparib once daily or CX-5461/olaparib combination for 3 weeks. The PDX were harvested at a tumor volume of 700 mm^3^. Mean tumor volume (mm^3^) ±95% CI (hashed lines represent individual mice) and corresponding Kaplan-Meier survival analysis. Censored events are represented by crosses on Kaplan-Meier plot. n indicates individual mice (4–8). **(C)** Gene expression heat maps utilising RNA expression data from the Broad Institute Cancer Cell Line Encyclopedia (CCLE) of the CX-5461 sensitivity gene signature encompassing BRCA network and MYC targets GSEA gene signatures identified in Figure 1E. The MYC signature (MYC_UP-sig) and BRCA-mutated (BRCAm-sig) signatures are enriched in the CX-5461-sensitive group compared to the resistant group *p*-value 0.004762 and 0.009524, respectively. **(D)** Clinical and RNA-seq gene expression data from 81 primary ovarian tumor samples from the Australian cohort of ovarian cancer patients from the International Cancer Genome Consortium (ICGC) (https://dcc.icgc.org/) (release 27). The level of expression of the CX-5461 sensitivity signatures were calculated using ssGSEA in individual samples. ssGSEA scores were normalized by linear transformation to the 0-1 range for comparison across primary and 25 ascites samples from relapse patients **(E)**. **(F)** Clustering of relapse samples with cell lines was based on the MYC_UP-sig and BRCAm-sig signatures of samples using k-means with a k=4. **(G)** Analysis of the MYC_UP-sig and BRCA-sig in three matching primary tumor samples of three relapsed samples in (E) enriched with CX-5461-sensitivity signature. **(H)** Clustering of samples in G with cell lines samples based on the MYC_UP-sig and BRCAm-sig signatures using k-means with a k=4.

To compare the *in vivo* efficacy of CX-5461 with standard-of-care therapy, we examined the activity of CX-5461 in chemo-naïve HGSOC-PDX (#62) with *BRCA1* promoter hypermethylation previously characterized as resistant/refractory to cisplatin and responsive to PARP inhibitor, rucaparib (37, 46). Interestingly, despite previously reported rucaparib response, PDX #62 did not respond to olaparib showing progressive disease with an increase in tumor volume of >20% from baseline at 8 days post-treatment (Figure 7B). In comparison, CX-5461 therapy led to stable disease and a statistically significant survival benefit (median TTH 92 days vs vehicle 32 days and olaparib 36 days, *p*-values 0.0004 and 0.0022 compared to vehicle and olaparib, respectively). The combination of CX-5461 and olaparib, however, provided no additional survival benefit. PDX#62 PDX harbours amplifications in multiple cancer-associated genes and increased expression of MYCN and Cyclin E (46), suggesting high oncogene-induced replication stress may confer strong sensitivity to CX-5461. This is consistent with the MYC targets gene expression signature being associated with sensitivity to CX-5461 in OVCA cell lines (Figure 1A&B). Importantly, our data clearly demonstrate that CX-5461 has a different mechanism of action and sensitivity spectrum to olaparib and is an exciting treatment option for patients that have poor response to platinum therapy (1).

In order to assess the potential of CX-5461 therapy in HGSOC treatment, we examined the prevalence of CX-5461 sensitivity gene expression signatures in HGSOC tumours samples. We built a generalized linear model using ssGSEA of the MYC (MYC-UP-Sig) and BRCA mutated (BRCAm-Sig) gene expression signatures identified in HGSOC cells lines as predictors for sensitivity to CX-5461 (Figure 1A & Figure 7C). We found that the model including both the BRCAm-sig and MYC_UP-sig signatures as predictors of sensitivity to CX-5461 provided a better fit than the model with the BRCAm-sig as sole predictor (ANOVA Chi-squared test p-value=0.026557). We then investigated the enrichment of these signatures in 81 primary ovarian tumor samples (Figure 7D) and 25 recurrent (ascites) samples (Figure 7E-F). 21 of the of the primary tumor samples (26%) exhibited the MYC_UP and BRCA-mutated gene expression signatures of sensitivity to CX-5461, of which 2 progressed while receiving chemotherapy and 13 relapsed following therapy (Figure 7D). In addition, 4 recurrent samples (16%) exhibit CX-5461 sensitivity gene signatures (Figure 7E-F). Intriguingly, the matching primary tumor of three of the relapsed tumors predicted to respond to CX-5461 did not exhibit the CX-5461 sensitivity gene signature. Thus, although these primary tumours are predicted to be resistant to CX-5461, upon the development of resistance to chemotherapy they acquired gene signatures of response to CX-5461 affirming our findings utilizing PDX#62 (Figure 7B) that CX-5461 is a treatment option for a selection of HGSOC with poor response to platinum and PARPi therapies.

## Discussion

### CX-5461 has significant therapeutic efficacy in HGSOC models

We have previously demonstrated that CX-5461 effectively treats MYC-driven lymphoma (19, 23) and, more recently AML independent of p53 status (20). CX-5461 activity is associated with activation of ATM/ATR signalling leading to S-phase delay and G2 cell cycle arrest (26, 27). CX-5461 was also shown to exhibit synthetic lethality with BRCA1/2 deficiency (25). While induction of the p53-dependent impaired ribosome biogenesis checkpoint is a major mechanism of efficacy of CX-5461 in p53-wild type tumors, activation of DDR is a key mechanism in the killing of p53-null AML and lymphoma (20, 27). As HGSOC is invariably p53-mutant and exhibits HR deficiency in up to 50% of cases, we hypothesized that targeted activation of DDR by CX-5461 would provide a novel therapeutic approach for HGSOC patients.

In this report, we demonstrate that CX-5461 has single agent therapeutic efficacy against HR-deficient HGSOC. Importantly, we demonstrate that CX-5461 has significant therapeutic efficacy against a cisplatin- and olaparib-resistant HGSOC PDX, demonstrating that CX-5461’s activity spectrum differs to that of cisplatin and olaparib. We have identified BRCA-mutated and MYC targets gene expression signatures as biomarkers for sensitivity to CX-5461. Our data reveal the BRCA mutation gene signature include defects in DNA repair pathways in addition to HR, such as BER that sensitize cancer cells to growth inhibition by CX-5461. We have identified predictive signatures of CX-5461 sensitivity in 26% of primary and 16% of relapsed ovarian cancer samples highlighting the potential of CX-5461 therapy in a subset of primary and acquired chemotherapy- and PARPi-resistant HGSOC. Specifically, we propose CX-5461 will have efficacy in HR-deficient HGSOC but also in HGSOC tumors with elevated MYC activity such as the high-MYCN HGSOC subtype associated with poor prognosis (34, 47).

### CX-5461 induces nucleolar DDR

Despite the fact that CX-5461 selectively inhibits Pol I transcription initiation (19, 42), its chronic effects on the stabilization of GQ structures have been proposed to impede the progression of DNA replication forks, rendering HR-deficient cancer cells sensitive to CX-5461 (25). Here, we demonstrate that CX-5461-mediated DDR is independent of its role in GQ stabilization, rather we show that by inhibiting Pol I transcription initiation, CX-5461 leads to R-loops stabilization, recruitment of RPA to ssDNA and ATR activation at the nucleoli in HR-proficient cells. Although we did not observe further increases in R-loops levels in HR-deficient cells, the levels of pRPA and pATR at UBF-bound rDNA regions at the periphery of the nucleoli were significantly increased, indicating the targeting of stalled rDNA replication forks to the periphery for repair (43). Thus, HRD potentiates CX-5461-mediated replication stress at rRNA genes highlighting compromised HR-dependent resolution of nucleolar replication stress as a potential mechanism of CX-5461 synthetic lethal interaction with HRD in HGSOC.

We demonstrate the net effect of CX-5461-induced nucleolar DDR is destabilization of replication forks via MRE11 activity leading to global replication stress and DNA damage. Recently, defects in stalled fork protection were identified as a common event (60%) in HGSOC patient-derived organoids (48). Therefore, CX-5461 may have efficacy in a subset of HGSOC with functional defects in replication fork protection. As CX-5461 activates DNA damage response and destabilizes replication forks irrespective of HR pathway status, it overcomes two well-known mechanisms of resistance to PARPi (HR restoration and stabilization of replication forks (12–15)).

### CX-5461 cooperates with PARPi in enhancing replication stress in HR-deficient HGSOC

The combination of CX-5461 and PARPi therapy showed robust therapeutic benefit in HR-deficient HGSOC, demonstrating that CX-5461’s interaction with PARPi can significantly improve treatment of HR-deficient HGSOC. CX-5461 combination with PARPi led to increased replication stress, DNA damage and cell death, consistent with their distinct mode of action in destabilizing replication forks and inducing replication stress. In the absence of BRCA and RAD51, nascent replication forks are extensively degraded by MRE11. Thus, we propose that CX-5461 exacerbates HRD-mediated degradation of replication forks leading to increased replication stress and accumulation of DNA damage. Therefore, the combined effect of CX-5461, PARPi and HRD in enhancing replication stress through differential effects on replication fork stability leads to the accumulation of DNA damage that underpins their strong cooperation in promoting cancer cell death.

### Clinical implications

Altogether, our data provide evidence for the potential of combining CX-5461 and PARPi for improving the treatment of HR-deficient HGSOC. We demonstrate that CX-5461 enhances the synthetic lethal interaction of PARPi with HRD and clearly show that CX-5461 has a different mechanism of action to PARPi. Importantly, we characterized BRCA-mutated and MYC targets gene signatures as predictors of patient’s response to CX-5461. As these predictive signatures were identified in primary and relapsed ovarian tumor samples, we propose that CX-5461 is an exciting treatment option not only for patients with HR-deficient tumors or tumors with unstable replication forks but also for patients with tumors that have high MYC activity and poor clinical outcome; these patients currently have very limited effective treatment options.

## Methods

### Compounds

BMN-673, olaparib, KU55933, VE-821, ABT-199 and everolimus were purchased from Selleckchem. CX-5461 was purchased from Synkinase.

### Cell lines

Individuality and the identity of ovarian cell lines listed in Supplementary Table 1 were routinely confirmed by a polymerase chain reaction (PCR) based short tandem repeat (STR) analysis using six STR loci. Generation of the WEHICS62 cell line from PDX #62 and the OVCAR8 RAD51C KO cell line is described in (37). Cell lines were maintained in culture for a maximum of 8-10 weeks. Mycoplasma testing was routinely performed by PCR.

### Cell proliferation assays

Cells were drug treated and cell number assessed via an imaging system (IncuCyte ZOOM) or the sulforhodamine B assay. To assess CX-5461 and BMN-673 anti-growth combination effects, dose response curves were generated for the single agents. Cells were less than 90% confluent in control wells at the end of incubation. GI doses were determined using GraphPad Prism.

For examining synthetic lethality of CX-5461 and siRNAs targeting the HR genes, individual siRNA duplexes (Dharmacon, GE lifesciences) were reversed transfected into OVCAR4 cells using Dharmafect 4 reagent (Dharmacon, GE lifesciences). 24h later, transfection medium was changed to either CX-5461 (80nM) or vehicle containing medium and cells were incubated further for 48h. Cell proliferation was measured by cell count using DAPI staining and imaging using Cellomics. Bliss Combination Index was calculated by dividing combined viability (siRNA plus CX-5461 treatment) by the multiply of individual viability (siRNA only or CX-5461 only treatment). Bliss Combination Index lower than 0.9 is considered as synergy. siRNAs Dharmacon catalogue No. siRAD54L (D-004592-17, D-004592-01, D-004592-02, D-004592-04); siRAD51AP1 (D-017166-01, D-017166-02, D-017166-03, D-017166-04); siBRCA2 (D-003462-04, D-003462-01, D-003462-02, D-003462-03).

### Cell cycle analysis

For cell cycle analysis using 5-bromo-2ʹ- deoxyuridine (BrdU) incorporation, cells were labelled with 10 μM BrdU for 30 min, washed twice with PBS, cells collected, pelleted and fixed in 80% ice-cold ethanol and stored at 4°C until further processing. Cells were pelleted and incubated in 1 mL of 2N HCl containing 0.5% (v/v) Triton X-100 for 30 min then pelleted and washed in 1 mL of 0.1M Na2B4O7.10H2O (pH 8.5). Cell pellets were sequentially incubated for 30 min with anti-BrdU and FITC anti-mouse IgG antibodies (Supplementary Table 2) in PBS containing 2% FBS and 0.5% Tween-20. Cells were washed with PBS- 2% FBS then incubated in 10 µg/mL propidium iodide (PI) solution at room temperature for 15 min. Cells were analyzed on the FACSCanto II (BD Biosciences) and cell cycle analysis was performed using Flowlogic software (Inivai Technologies).

### Cell death assay

Cell death was determined using PI staining followed by flow cytometry (FACSCanto II) and data analyzed using Flowlogic software.

### Clonogenic assays

Cells were seeded in 6-well plates for 24h. Following drug incubation for 5 days, media was aspirated, and cells were washed and incubated with drug-free media for 7 days. Cells were fixed with 100% methanol for 1h, stained with 0.1% (w/v) crystal violet for 1h, washed with H_2_O and air dried. Colonies was counted manually with a stereomicroscope.

### Gene expression analysis

Cells were harvested at 50–80% confluency (3 biological replicates). RNA was extracted (QIAGEN RNeasy kit), *in vitro* transcribed and biotin labelled cRNA was fragmented and hybridized to Affymetrix 1.0ST expression array as per manufacturer’s instructions. Differential gene expression was determined using the Limma R package after RMA normalisation and back-ground correction (49). Genes that had a >1.4-fold change in expression between resistant and sensitive were included in the MetaCore pathway analysis (http://thomsonreuters.com/metacore).

For GSEA 1000 iterations were performed using the default weighted enrichment statistic and a signal-to-noise metric to rank genes based on their differential expression across sensitive and resistant cell lines (50).

We used ssGSEA (32) from the GSVA (33) package (version 1.20.0) in R (version 3.3.2) to obtain the level of activity of pathways utilizing GeneGo (MetaCore) gene ontologies for the HR, BER and NHEJ pathways and the HRD gene signature reported previously (30) in individual samples. Here, genes in each sample were ranked according to their expression levels, and a score for each pathway was generated based on the empirical cumulative distribution function, reflecting how highly or lowly genes of a pathway are found in the ranked list. Statistical significance of the ssGSEA scores of different cell line categories (sensitive or resistant) was obtained using two-sided Wilcoxon tests. Benjamini-Hochberg correction was subsequently used to account for multiple testing.

For Reverse-transcription qPCR analysis, cells were lysed, RNA was extracted, and first-strand cDNA was synthesized using random hexamer primers and Superscript III (Invitrogen). Quantitative PCR (qPCR) was performed in duplicate using the FAST SYBR Green dye on the StepOnePlus real-time PCR system (Applied Biosystems). Primer sequences are listed in (Supplementary Table 3).

### Western blotting (WB)

Twenty to fifty micrograms of whole-cell lysates were resolved by SDS-PAGE, electrophoretically transferred onto PVDF membranes (Milli-pore) and analyzed using enhanced chemiluminescence (ECL) detection (GE Healthcare). Antibodies details are listed in Supplementary Table 3.

### Immunofluorescence

For IF assays combined with EdU labelling, cells were first incubated with 10 μM EdU for 30 min prior to drug treatment. Cells were fixed in 4% paraformaldehyde (PFA) (10 min at room temperature), permeabilized with 0.3% Triton X-100 in PBS for 10 min on ice, washed with PBS, and blocked with 5% goat serum and 0.3% Triton X-100 in PBS for 30 min. Cells were sequentially incubated with the primary antibody and secondary antibodies (Supplementary Table 2). Cells were incubated for 30 min at room temperature in Click-IT reaction (100mM Tris pH 8.5, 10 nM Alexa Fluor 647-azide (Cat# A10277, Thermo Fisher Scientific), 1 mM CuSO4, and 100mM ascorbic acid), then washed with PBS. Nuclei were counterstained with DAPI in Vectashield mounting media (Vector Labs).

For IF using the 1H6 antibody for detection of GQ DNA, cells were treated with 40 μg/ml RNase A for 1h prior to the blocking step. For IF using the S9.6 (R-loops) antibody, cells post-PFA fixation were permeabilized with 100% methanol for 10 min and 100% acetone for 1min on ice, washed with PBS prior to the blocking step. Images were acquired on an Olympus BX-61 microscope equipped with a Spot RT camera (model 25.4), using the UPlanAPO 60X, NA 1.2 water immersion objective and the Spot Advanced software, version 4.6.4.3. Settings for adjusting the image after acquisition (i.e. gamma adjust and background subtract settings) were identical for all images. Confocal images were acquired using Zeiss Elyra 63X magnification. Images were analyzed using Cell Profiler.

### DNA fibre analysis

Exponentially growing OVCAR8 cells were pulse-labelled with 50 μM CldU (20 min), washed and exposed to 250 μM IdU (20 min). After exposure to the second nucleotide analog, the cells were washed again in warm 1X PBS and either processed or treated for 3 hr with 1 μM CX-5461, mirin (50 μM, Sigma-Aldrich) or CX-5461 + mirin. Labelled cells were trypsinized and resuspended in ice-cold PBS at 7.5 × 10^5^ cells/mL. Two microliters of this suspension were spotted onto a pre-cleaned glass slide and lysed with 10 μL of spreading buffer (0.5% SDS in 200 mM Tris-HCl, pH 7.4 and 50 mM EDTA) in a humid chamber. After 36 min, the slides were tilted at 15° relative to horizontal, allowing the DNA to spread. Slides were air-dried, fixed in methanol and acetic acid (3:1) for 10 min and air-dried. DNA was denatured with 2.5 M HCl for 60 min at room temperature. Slides were then rinsed in PBS thrice and blocked in PBS + 0.1% Triton X-100 (PBS-T) + 1% BSA for 1 hr at room temperature. Rat anti-BrdU (1:200, Abcam ab6323) was applied overnight at 4°C in a humid chamber. Slides were then washed with PBS and incubated with Alexa Fluor 488-conjugated chicken anti-rat antibody at 1:200 dilutions (Life technologies, A21470). Slides were washed wit PBS and incubated for 45 minutes at room temperature with mouse anti-BrdU (Becton Dickinson, 347580) antibody at 1:50 dilution to detect IdU tracks. Slides were washed in PBS and stained with Alexa Fluor 594-labelled goat anti-mouse antibody (Life technologies, A-11030) at 1:300 dilutions at room temperature for 30 min. Slides were washed in PBS and mounted in Prolong Diamond antifade (Invitrogen, P36961). Replication tracks were imaged on a Deltavision microscope at 60X and measured using ImageJ software. In each experiment, 150 or more individual tracks were measured for fork degradation estimation. Data are representative of at least two independent experiments. For studies with CX5461 and BMN-673, the cells were treated for 24 hours with individual drugs or in combination before cells were labelled with the BrdU analogs.

### Animal studies

All experiments involving animals were approved by the Walter and Eliza Hall Institute of Medical Research Animal Ethics Committee. PDX were generated from with patients with OVCA enrolled in the Australian Ovarian Cancer Study. Additional ethics approval was obtained from the Human Research Ethics Committees at the Royal Women’s Hospital and the Walter and Eliza Hall Institute.

PDX were generated as published previously by transplanting fresh fragments subcutaneously into NOD/SCID/IL2Rγnull recipient mice (T1, passage 1) (46). Briefly, for PDX #19B fresh tumor fragments were subcutaneously implanted under the right flank. For PDX #62 frozen tumor fragments from previously passaged PDX (46), stored in DMSO supplemented media, were thawed and subcutaneously implanted under the right flank. Recipient mice bearing T4-T7 (passage 4 to passage 7) tumors were randomly assigned to treatment with olaparib, CX-5461, combination or vehicle when tumor volume reached 180-300 mm^3^. Olaparib was administered once daily intraperitoneally at a dose of 50 mg/kg in vehicle (phosphate buffered saline (PBS) containing 10% DMSO and 10% 2-hydroxy-propyl-β-cyclodextrin). CX-5461 was given by oral gavage twice a week for 3 weeks at 40 mg/kg in vehicle (25mM Na H_2_ PO_4_ pH7.4). Tumors were measured twice per week and recorded in StudyLog software (StudyLog Systems). Mice were euthanized once tumor volume reached 700 mm^3^ or when mice reached ethical endpoint. Nadir, time to harvest (TTH) and treatment responses are as defined previously (46). Tumor volume and survival graphs were produced with SurvivalVolume v1.2 (51). Median TTH was calculated by including censored events for PDX where mice were harvested when tumor volume was > 500 mm^3^ but < 600 mm^3^. Partial response was achieved if the average tumor volume reduced to between 50 mm^3^ and 140 mm^3^ (> 30% reduction from nadir, assigned as 200 mm^3^) for two or more consecutive measurements.

### Analysis of ovarian tumor samples

We obtained clinical and RNAseq gene expression data from 81 primary solid tumour samples and 25 ascites samples from relapse patients from the Australian cohort of ovarian cancer patients available from the International Cancer Genome Consortium (https://dcc.icgc.org/) (release 27). Only coding genes were considered. We calculated the level of expression of the MYC_UP (MYC oncogenic Signature UP) and BRCAm (BRCA1 Mutated UP) signatures using ssGSEA in individual samples. ssGSEA scores were normalized by linear transformation to the 0-1 range for comparison across primary, relapse, and cell line data.

Clustering of relapse samples with cell lines was based on the BRCAm and MYC_UP signatures of samples using k-means with a k=4.

## Abbreviations

ATM: Ataxia telangiectasia mutated
ATR: Ataxia telangiectasia and Rad3
BER: Base excision repair
CHK1/2: Checkpoint kinases 1/2
DDR: DNA damage response
DSBs: double strand breaks
DDR therapy: DNA repair and DDR inhibitors
5’ETS: External spliced region
GI: Growth inhibition
GQ: G-quadruplex DNA
HGSOC: High-grade serous ovarian cancer
HR: homologous recombination
HRD: HR deficiency
IR: ionizing radiation
KO: Knockout
TTH: time to harvest
NHEJ: Non-homologous end joining
OVCA: Ovarian cancer
PDX: Patient-derived xenografts
PARP: Poly-(ADP-ribose) polymerase
PARPi: PARP inhibitors
Pol I: RNA polymerase I
RPA: replication protein A
rRNA: ribosomal RNA
rDNA: rRNA genes
R-loops: RNA DNA hybrids
ssDNA: single-strand DNA
TSS: Transcription start site
UBF: upstream binding transcription factor

## Authors’ Contributions

Conception and design: E. Sanij, K. E. Sheppard, R.B. Pearson

Development of methodology: E. Sanij, K. Hannan, S. Yan, J. Xuan, K.T. Chan, J. Kang, S. Ellis, C. Cullinane, M. Wakefield, E. Berns

Acquisition of data: E. Sanij, K. Hannan, J. Xuan, J. Ahern, J. Son, O. Kondrashova, E. Lieschke, P. Nag, D. Frank

Analysis and interpretation of data (e.g., statistical analysis, computational analysis): E. Sanij, A. Trigos, J. Kang, K. Khanna, G. Poortinga, L. Mileshkin, G. McArthur, J. Soong, R.D. Hannan, C. Scott, K. Sheppard, R.B. Pearson

Writing, review, and/or revision of the manuscript: E. Sanij, K. Hannan, C. Cullinane, G. Poortinga, K. Khanna, R.D. Hannan, C. Scott, K. Sheppard, R.B. Pearson Study supervision: E. Sanij, K. Sheppard, R.B. Pearson

## References

1. Cancer Genome Atlas Research N. Integrated genomic analyses of ovarian carcinoma. Nature. 2011;474(7353):609–15.

2. Ahmed AA, Etemadmoghadam D, Temple J, Lynch AG, Riad M, Sharma R, et al. Driver mutations in TP53 are ubiquitous in high grade serous carcinoma of the ovary. J Pathol. 2010;221(1):49–56.

3. Konstantinopoulos PA, Ceccaldi R, Shapiro GI, and D’Andrea AD. Homologous Recombination Deficiency: Exploiting the Fundamental Vulnerability of Ovarian Cancer. Cancer Discov. 2015;5(11):1137–54.

4. Lord CJ, and Ashworth A. PARP inhibitors: Synthetic lethality in the clinic. Science. 2017;355(6330):1152–8.

5. Scott CL, Swisher EM, and Kaufmann SH. Poly (ADP-ribose) polymerase inhibitors: recent advances and future development. J Clin Oncol. 2015;33(12):1397–406.

6. Maya-Mendoza A, Moudry P, Merchut-Maya JM, Lee M, Strauss R, and Bartek J. High speed of fork progression induces DNA replication stress and genomic instability. Nature. 2018;559(7713):279–84.

7. Ceccaldi R, Liu JC, Amunugama R, Hajdu I, Primack B, Petalcorin MI, et al. Homologous-recombination-deficient tumours are dependent on Poltheta-mediated repair. Nature. 2015;518(7538):258–62.

8. Swisher EM, Lin KK, Oza AM, Scott CL, Giordano H, Sun J, et al. Rucaparib in relapsed, platinum-sensitive high-grade ovarian carcinoma (ARIEL2 Part 1): an international, multicentre, open-label, phase 2 trial. Lancet Oncol. 2017;18(1):75–87.

9. Drean A, Lord CJ, and Ashworth A. PARP inhibitor combination therapy. Crit Rev Oncol Hematol. 2016;108:73–85.

10. Mariappan L, Jiang XY, Jackson J, and Drew Y. Emerging treatment options for ovarian cancer: focus on rucaparib. Int J Womens Health. 2017;9:913–24.

11. Moore K, Colombo N, Scambia G, Kim BG, Oaknin A, Friedlander M, et al. Maintenance Olaparib in Patients with Newly Diagnosed Advanced Ovarian Cancer. N Engl J Med. 2018.

12. Barber LJ, Sandhu S, Chen L, Campbell J, Kozarewa I, Fenwick K, et al. Secondary mutations in BRCA2 associated with clinical resistance to a PARP inhibitor. J Pathol. 2013;229(3):422–9.

13. Edwards SL, Brough R, Lord CJ, Natrajan R, Vatcheva R, Levine DA, et al. Resistance to therapy caused by intragenic deletion in BRCA2. Nature. 2008;451(7182):1111–5.

14. Kondrashova O, Nguyen M, Shield-Artin K, Tinker AV, Teng NNH, Harrell MI, et al. Secondary Somatic Mutations Restoring RAD51C and RAD51D Associated with Acquired Resistance to the PARP Inhibitor Rucaparib in High-Grade Ovarian Carcinoma. Cancer Discov. 2017;7(9):984–98.

15. Chaudhuri AR, Callen E, Ding X, Gogola E, Duarte AA, Lee JE, et al. Erratum: Replication fork stability confers chemoresistance in BRCA-deficient cells. Nature. 2016;539(7629):456.

16. Diesch J, Hannan RD, and Sanij E. Perturbations at the ribosomal genes loci are at the centre of cellular dysfunction and human disease. Cell Biosci. 2014;4:43.

17. Hein N, Hannan KM, George AJ, Sanij E, and Hannan RD. The nucleolus: an emerging target for cancer therapy. Trends Mol Med. 2013;19(11):643–54.

18. Pelletier J, Thomas G, and Volarevic S. Ribosome biogenesis in cancer: new players and therapeutic avenues. Nat Rev Cancer. 2018;18(1):51–63.

19. Bywater MJ, Poortinga G, Sanij E, Hein N, Peck A, Cullinane C, et al. Inhibition of RNA polymerase I as a therapeutic strategy to promote cancer-specific activation of p53. Cancer Cell. 2012;22(1):51–65.

20. Hein N, Cameron DP, Hannan KM, Nguyen NN, Fong CY, Sornkom J, et al. Inhibition of Pol I transcription treats murine and human AML by targeting the leukemia-initiating cell population. Blood. 2017.

21. Peltonen K, Colis L, Liu H, Jäämaa S, Zhang Z, Af Hällström T, et al. Small molecule BMH-compounds that inhibit RNA polymerase I and cause nucleolar stress. Mol Cancer Ther. 2014;13(11):2537–46.

22. Peltonen K, Colis L, Liu H, Trivedi R, Moubarek MS, Moore HM, et al. A targeting modality for destruction of RNA polymerase I that possesses anticancer activity. Cancer Cell. 2014;25(1):77–90.

23. Devlin JR, Hannan KM, Hein N, Cullinane C, Kusnadi E, Ng PY, et al. Combination Therapy Targeting Ribosome Biogenesis and mRNA Translation Synergistically Extends Survival in MYC-Driven Lymphoma. Cancer Discov. 2016;6(1):59–70.

24. Khot A, Brajanovski N, Cameron D, Poortinga G, Sanij E, Lim J, et al. American Society of Heamatology. Atlanta, GA, USA; 2017.

25. Xu H, Di Antonio M, McKinney S, Mathew V, Ho B, O’Neil NJ, et al. CX-5461 is a DNA G-quadruplex stabilizer with selective lethality in BRCA1/2 deficient tumours. Nat Commun. 2017;8:14432.

26. Negi SS, and Brown P. rRNA synthesis inhibitor, CX-5461, activates ATM/ATR pathway in acute lymphoblastic leukemia, arrests cells in G2 phase and induces apoptosis. Oncotarget. 2015;6(20):18094–104.

27. Quin J, Chan KT, Devlin JR, Cameron DP, Diesch J, Cullinane C, et al. Inhibition of RNA polymerase I transcription initiation by CX-5461 activates non-canonical ATM/ATR signaling. Oncotarget. 2016.

28. Cornelison R, Dobbin ZC, Katre AA, Jeong DH, Zhang Y, Chen D, et al. Targeting RNA-Polymerase I in Both Chemosensitive and Chemoresistant Populations in Epithelial Ovarian Cancer. Clin Cancer Res. 2017;23(21):6529–40.

29. Sheppard KE, Cullinane C, Hannan KM, Wall M, Chan J, Barber F, et al. Synergistic inhibition of ovarian cancer cell growth by combining selective PI3K/mTOR and RAS/ERK pathway inhibitors. Eur J Cancer. 2013;49(18):3936–44.

30. Peng G, Chun-Jen Lin C, Mo W, Dai H, Park YY, Kim SM, et al. Genome-wide transcriptome profiling of homologous recombination DNA repair. Nat Commun. 2014;5:3361.

31. Pitroda SP, Pashtan IM, Logan HL, Budke B, Darga TE, Weichselbaum RR, et al. DNA repair pathway gene expression score correlates with repair proficiency and tumor sensitivity to chemotherapy. Sci Transl Med. 2014;6(229):229ra42.

32. Barbie DA, Tamayo P, Boehm JS, Kim SY, Moody SE, Dunn IF, et al. Systematic RNA interference reveals that oncogenic KRAS-driven cancers require TBK1. Nature. 2009;462(7269):108–12.

33. Hanzelmann S, Castelo R, and Guinney J. GSVA: gene set variation analysis for microarray and RNA-seq data. BMC Bioinformatics. 2013;14:7.

34. Jung M, Russell AJ, Liu B, George J, Liu PY, Liu T, et al. A Myc Activity Signature Predicts Poor Clinical Outcomes in Myc-Associated Cancers. Cancer Res. 2017;77(4):971–81.

35. Tothill RW, Tinker AV, George J, Brown R, Fox SB, Lade S, et al. Novel molecular subtypes of serous and endometrioid ovarian cancer linked to clinical outcome. Clin Cancer Res. 2008;14(16):5198–208.

36. Domcke S, Sinha R, Levine DA, Sander C, and Schultz N. Evaluating cell lines as tumour models by comparison of genomic profiles. Nat Commun. 2013;4:2126.

37. Kondrashova O, Topp M, Nesic K, Lieschke E, Ho GY, Harrell MI, et al. Methylation of all BRCA1 copies predicts response to the PARP inhibitor rucaparib in ovarian carcinoma. Nat Commun. 2018;9(1):3970.

38. Sirbu BM, Couch FB, Feigerle JT, Bhaskara S, Hiebert SW, and Cortez D. Analysis of protein dynamics at active, stalled, and collapsed replication forks. Genes Dev. 2011;25(12):1320–7.

39. Santos-Pereira JM, and Aguilera A. R loops: new modulators of genome dynamics and function. Nat Rev Genet. 2015;16(10):583–97.

40. Sanij E, Diesch J, Lesmana A, Poortinga G, Hein N, Lidgerwood G, et al. A novel role for the Pol I transcription factor UBTF in maintaining genome stability through the regulation of highly transcribed Pol II genes. Genome Res. 2015;25(2):201–12.

41. Sanij E, Poortinga G, Sharkey K, Hung S, Holloway TP, Quin J, et al. UBF levels determine the number of active ribosomal RNA genes in mammals. The Journal of cell biology. 2008;183(7):1259–74.

42. Drygin D, Lin A, Bliesath J, Ho CB, O’Brien SE, Proffitt C, et al. Targeting RNA polymerase I with an oral small molecule CX-5461 inhibits ribosomal RNA synthesis and solid tumor growth. Cancer Res. 2011;71(4):1418–30.

43. van Sluis M, and McStay B. A localized nucleolar DNA damage response facilitates recruitment of the homology-directed repair machinery independent of cell cycle stage. Genes Dev. 2015;29(11):1151–63.

44. Taglialatela A, Alvarez S, Leuzzi G, Sannino V, Ranjha L, Huang JW, et al. Restoration of Replication Fork Stability in BRCA1- and BRCA2-Deficient Cells by Inactivation of SNF2-Family Fork Remodelers. Mol Cell. 2017;68(2):414–30 e8.

45. Guetg C, Scheifele F, Rosenthal F, Hottiger MO, and Santoro R. Inheritance of silent rDNA chromatin is mediated by PARP1 via noncoding RNA. Mol Cell. 2012;45(6):790–800.

46. Topp MD, Hartley L, Cook M, Heong V, Boehm E, McShane L, et al. Molecular correlates of platinum response in human high-grade serous ovarian cancer patient-derived xenografts. Mol Oncol. 2014;8(3):656–68.

47. Helland A, Anglesio MS, George J, Cowin PA, Johnstone CN, House CM, et al. Deregulation of MYCN, LIN28B and LET7 in a molecular subtype of aggressive high-grade serous ovarian cancers. PLoS One. 2011;6(4):e18064.

48. Hill SJ, Decker B, Roberts EA, Horowitz NS, Muto MG, Worley MJ, Jr., et al. Prediction of DNA Repair Inhibitor Response in Short-Term Patient-Derived Ovarian Cancer Organoids. Cancer Discov. 2018;8(11):1404–21.

49. Smyth GK, Michaud J, and Scott HS. Use of within-array replicate spots for assessing differential expression in microarray experiments. Bioinformatics. 2005;21(9):2067–75.

50. Subramanian A, Tamayo P, Mootha VK, Mukherjee S, Ebert BL, Gillette MA, et al. Gene set enrichment analysis: a knowledge-based approach for interpreting genome-wide expression profiles. Proc Natl Acad Sci U S A. 2005;102(43):15545–50.

51. Wakefield MJ. survivalvolume, https://github.com/genomematt/survivalvolume. Github repository. 2016.

