## Supplemental data for "Inhibition of RNA Polymerase I Transcription Activates Targeted DNA Damage Response and Enhances the Efficacy of PARP Inhibitors in High-Grade Serous Ovarian Cancer"

Supplementary Data
Manuscript: Sanij et al.,

Title: **Inhibition of RNA Polymerase I Transcription Activates Targeted DNA Damage Response and Enhances the Efficacy of PARP Inhibitors in High-Grade Serous Ovarian Cancer**

**Content:**
Supplementary Figures 1-7.
Supplementary Tables 1-3.

#### Supplementary Figure 1

##### PUJANA_BRCA1_Centered_Network_GSEA

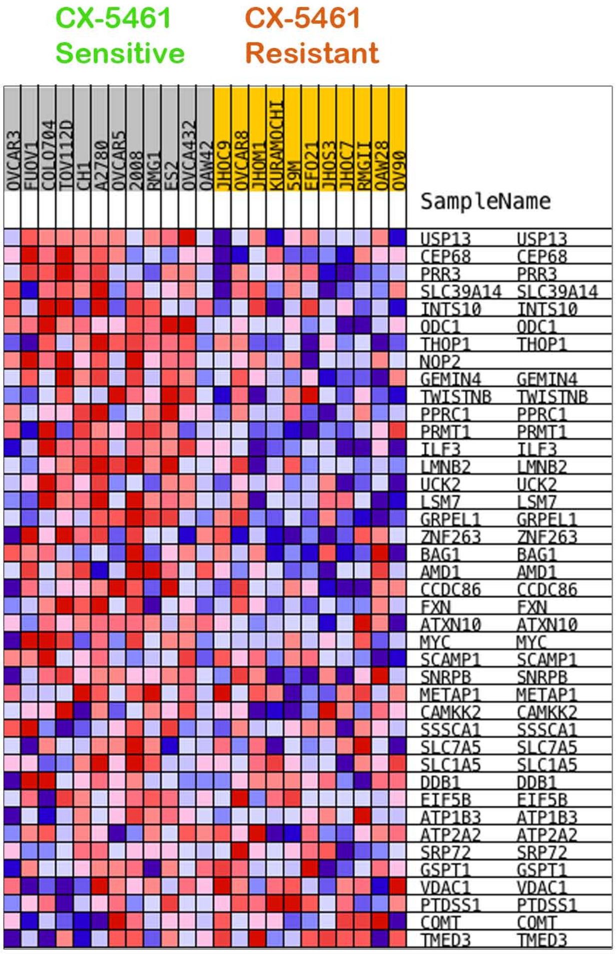


##### SCHLOSSER_ MYC_Targets_And_Serum_Response_UP_GSEA

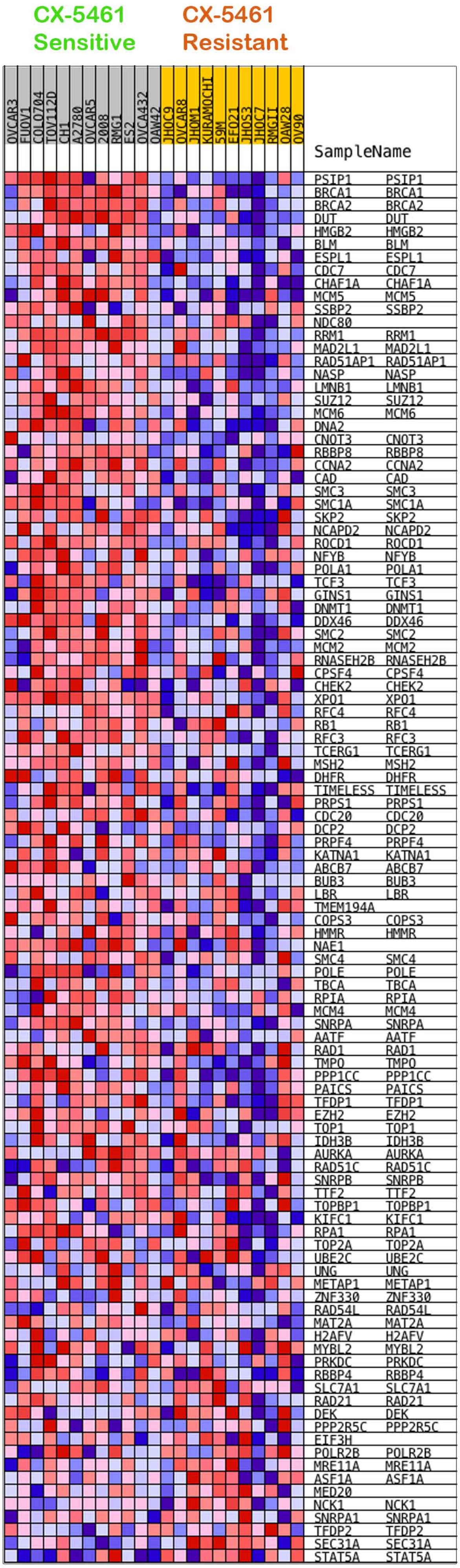

**Supplementary Figure 1**

Gene set enrichment analysis of microarray expression data of 12 CX-5461-sensitive and 11 -resistant cell lines. The analysis identified that the BRCA network and MYC targets GSEA gene sets to be enriched in CX-5461-sensitive OVCA cell lines. The heatmaps demonstrate relatively high (red) or low (blue) gene expression in the indicated sample.

**Supplementary Figure 2**

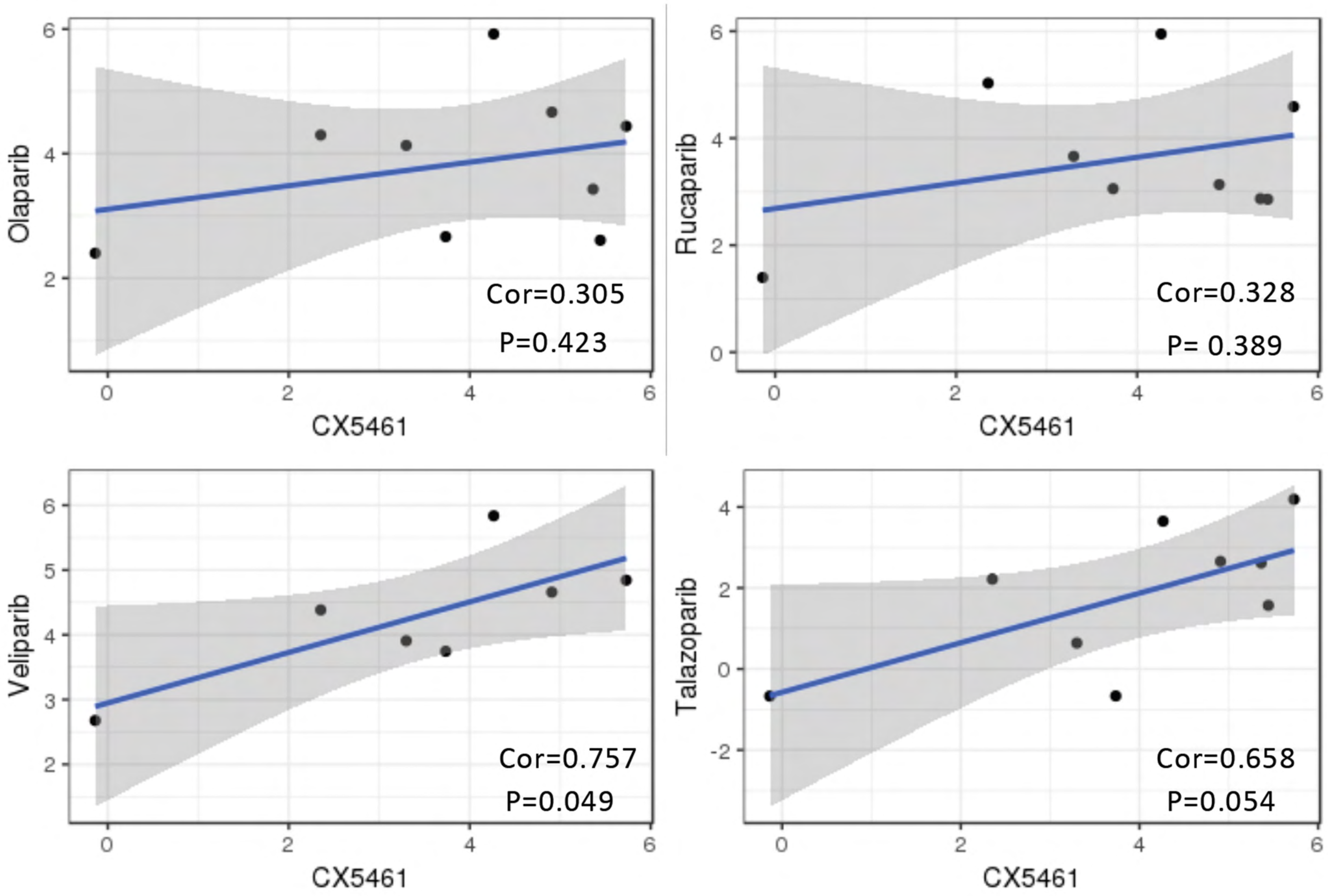

**Supplementary Figure 2**

Correlation between drug sensitivity measurements of CX-5461 and various PARPi in OVCA cell lines obtained from the Genomics of Drug Sensitivity database. Although a correlation between the sensitivity profiles of CX-5461 and PARPi ranging from 0.3 to 0.7 was observed, this was found to not be significant in 3 (Olaparib, Rucaparib and Talazoparib) out of the 4 PARPi investigated.

### Supplementary Figure 3

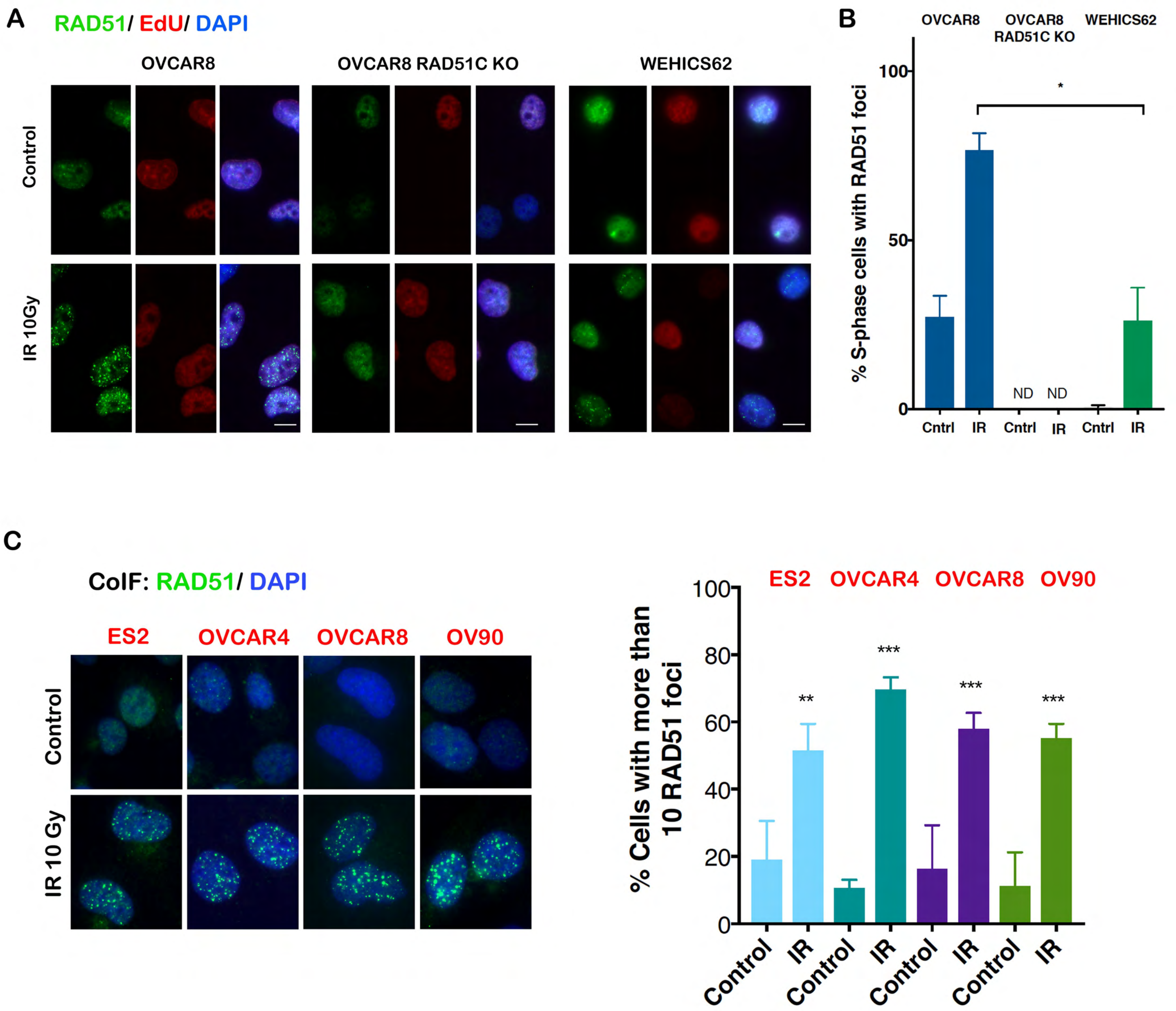

**Supplementary Figure 3.** Assessment of HR proficiency in OVCA cell lines. **A)** Assessment of HR proficiency in HGSOC cell lines. RAD51 foci formation assessed 6 hours post exposure to 10 Gy IR irradiation in HR-competent HGSOC cell line (OVCAR8), OVCAR8 derivative with RAD51C KO, and patient-derived HGSOC cell line with homozygous *BRCA1* hypermethylation (WEHICS62). Cells were treated with 10 μM EdU then irradiated and incubated for 6 hours. Cells were fixed with 10% paraformaldehyde and immunofluorescence for RAD51 was performed. Cells were incubated for 30 minutes at room temperature in Click-IT reaction (100 mM Tris pH 8.5, 10 nM Alexa Fluor 647-azide (Cat# A10277, Thermo Fisher Scientific), 1mM $CuSO_4$ and 100 mM Ascorbic Acid) then washed with PBS and counterstained with DAPI. **B)** Quantification of S-phase (EdU-positive) cells exhibiting > 10 RAD51 foci per cell. Mean ± SEM, $n$= 3, >170 EdU-positive cells were counted per condition. Statistical analysis was performed using student t test, $*p < 0.05$. ND, denotes not detected. **C)** RAD51 foci formation assessed 6 hours post exposure to 10 Gy IR irradiation. Quantification of cells exhibiting > 10 RAD51 foci per cell (right panel). Mean ±SEM, $n$= 3, >250 cells were counted per condition. Statistical analysis was performed using one-way ANOVA multiple comparisons, $*p < 0.05$, $**p < 0.01$, $***p$-value < 0.001.

#### Supplementary Figure 4

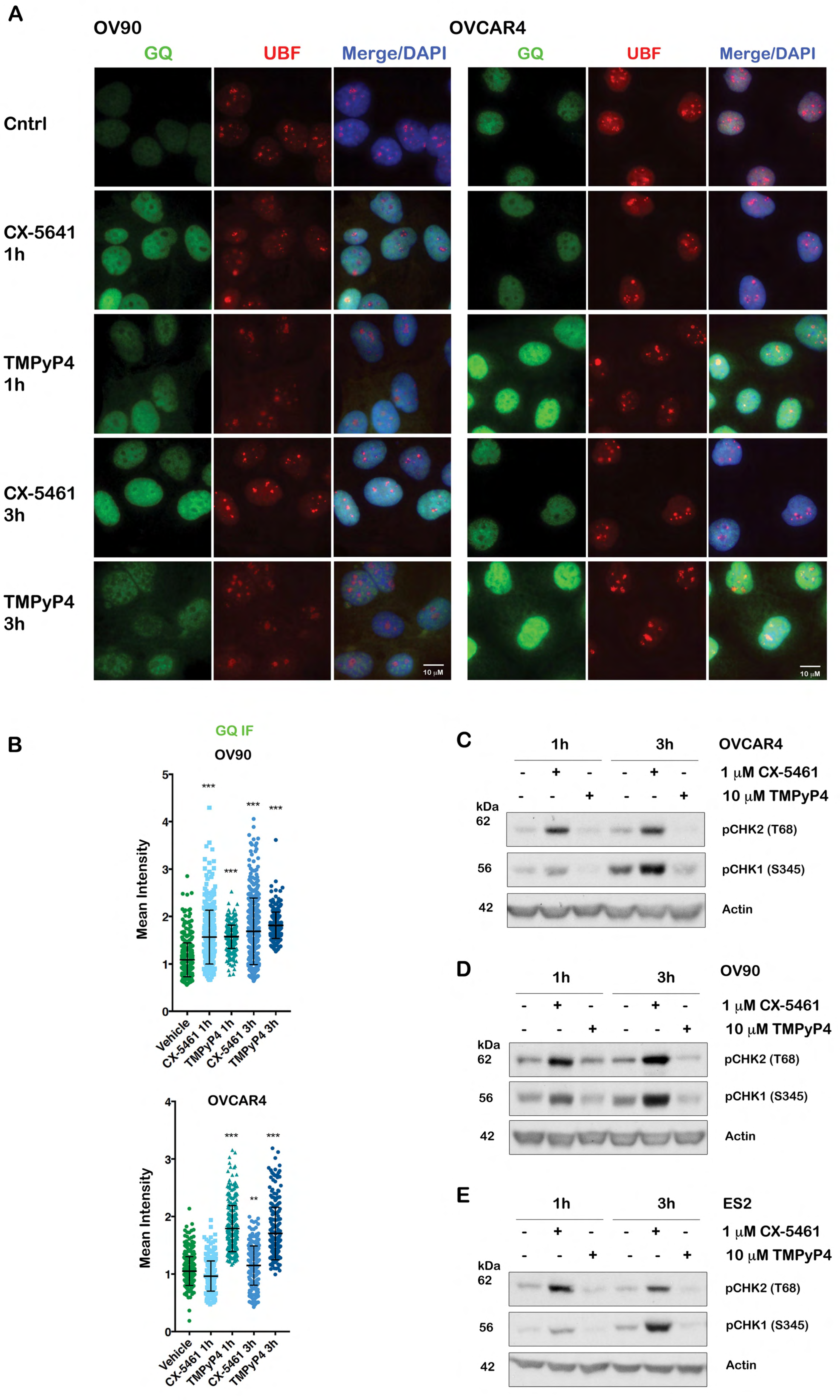

**Supplementary Figure 4**

CX-5461 induces DDR independent of its ability to stabilise GQ DNA structures. **A)** Co-immunofluorescence (Co-IF) of GQ DNA and UBF as a nucleolar marker in OV90 and OVCAR4 cells treated with either vehicle, 1 μM CX-5461 or 10 μM TMPyP4 as indicated. Representative images of *n* =2 for OV90 and *n* =3 for OVCAR4. **B)** Quantitation of GQ DNA immunofluorescence. Signal intensities were analyzed using Cell Profiler and normalized to corresponding vehicle controls. Error bars represent mean ± SD, OV90 *n* =2, >345 cells per condition, OVCAR4 *n* =3, >240 cells per condition. Statistical analysis of the increase in GQ signal intensity was performed using Kruskal-Wallis test, $^{**}p$-value < 0.01, $^{***}p$-value < 0.001 compared to vehicle control. **C-E)** CX-5461, but not TMPyP4, induces DDR. Cells were treated with vehicle, CX-5461 or TMPyP4 for 1h or 3h. Total protein lysates were analysed by western blotting. Representative blots of *n* =3.

### Supplementary Figure 5

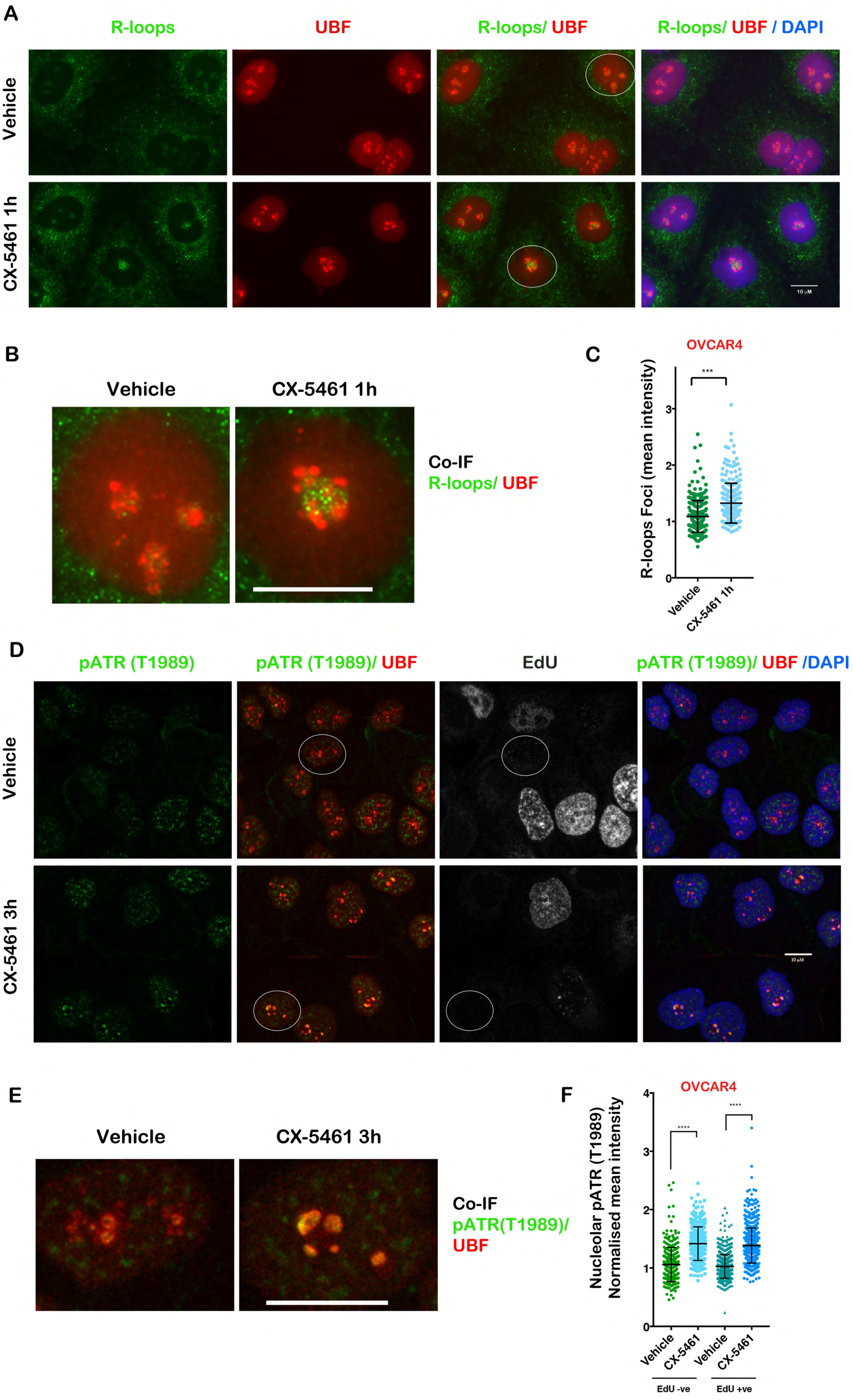

**Supplementary Figure 5**

**A)** Co-IF analysis of R-loops and UBF in OVCAR4 cells treated with vehicle or 1 μM CX-5461 for 3h. Cells emphasized by white circles are enlarged in (**B**). Scale bare denotes 10 μM. **C)** Quantitation of R-loops signal intensity of assays as in **A** was performed using Cell Profiler and normalized to the median of vehicle treated controls. $n$ =3, >260 cells per condition. Statistical analysis was performed using Mann-Whitney test, ***$p$-value < 0.001. **D)** Co-IF analysis of pATR(T1989) and UBF in OVCAR4 cells labeled with EdU and treated with vehicle 1 μM CX-5461 for 3h. EdU -ve cells emphasized by white circles are enlarged in (**E**). Scale bare denotes 10 μM. **F)** Quantitation of signal intensity of the colocalized regions was performed using Cell Profiler and normalized to the median of vehicle treated controls. Statistical analysis was performed using Mann-Whitney test, $n$ =3, >450 cells per condition, ***$p$-value < 0.001.

### Supplementary Figure 6

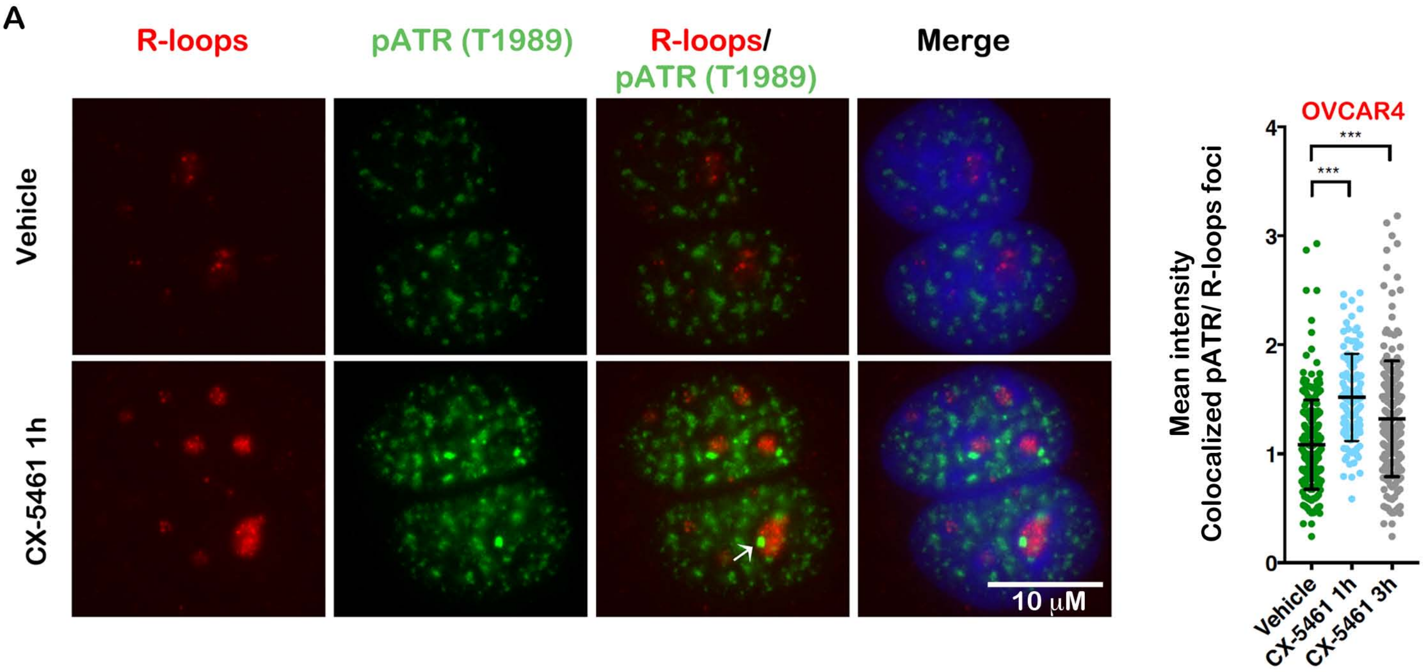


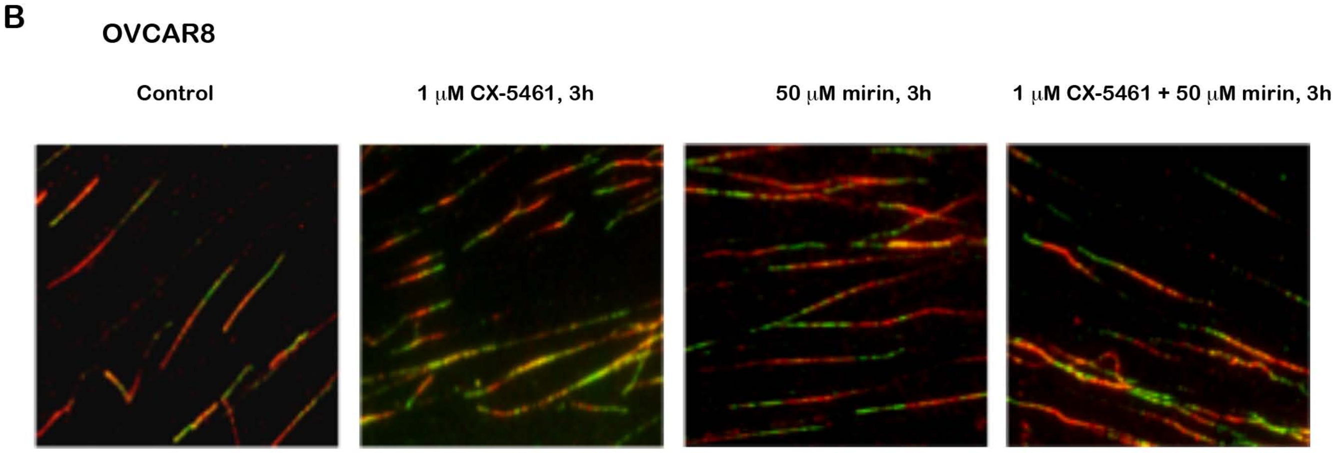

**Supplementary Figure 6.**

**A)** Co-immunofluorescence analysis of R-loops and pATR(T1989) in OVCAR4 cells treated with vehicle or 1 µM CX-5461 for 1h, n=1 or 3h n=2 >100 cells per condition. Signal intensity was normalized to median vehicle control. Error bars represent mean ± SD. Statistical analysis was performed using the Kruskal-Wallis test, ***p-value < 0.001 compared to vehicle control. **B)** Representative images of DNA fibre analysis of experiments presented in Figure 5D. OVCAR8 cells were sequentially labelled and either processed or treated with 1 µM CX-5461, 50 mM mirin or CX-5461+ mirin for 3h. Fibres were processed for DNA fibre analysis; *n*=2. Replication Fork length was calculated based on the length of the IdU tracks measured using ImageJ software. At least 150 replication tracks were analysed per experiment.

#### Supplementary Figure 7

A

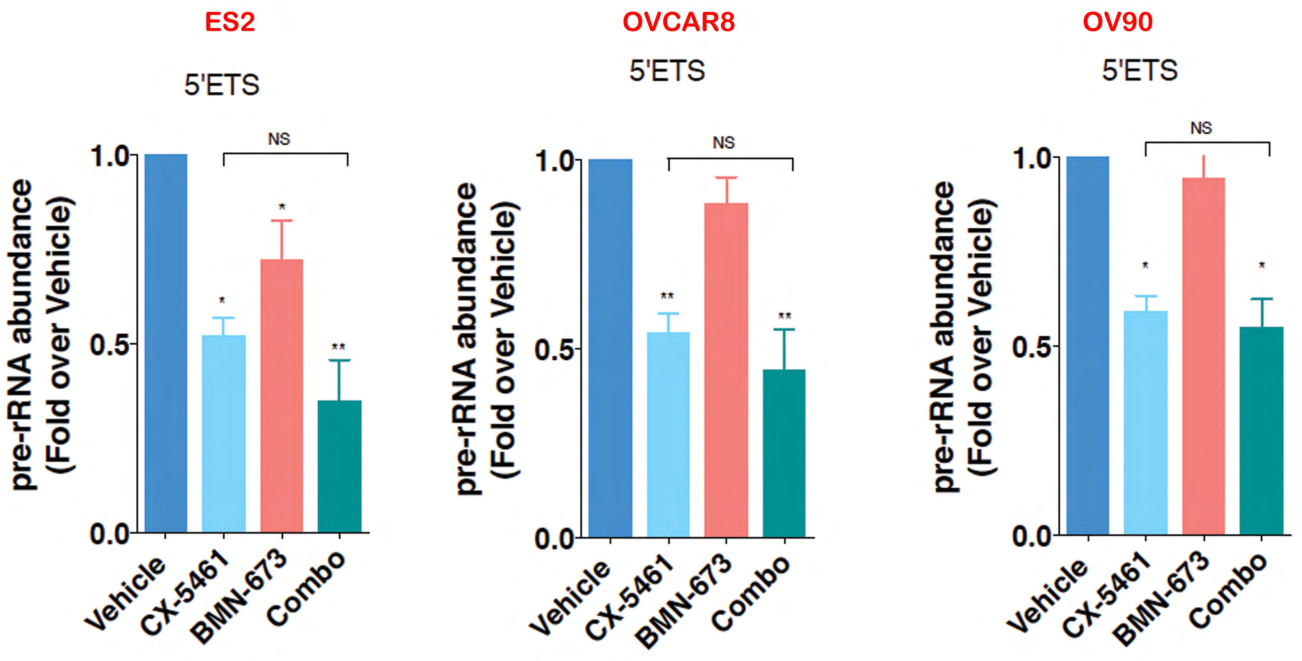


B

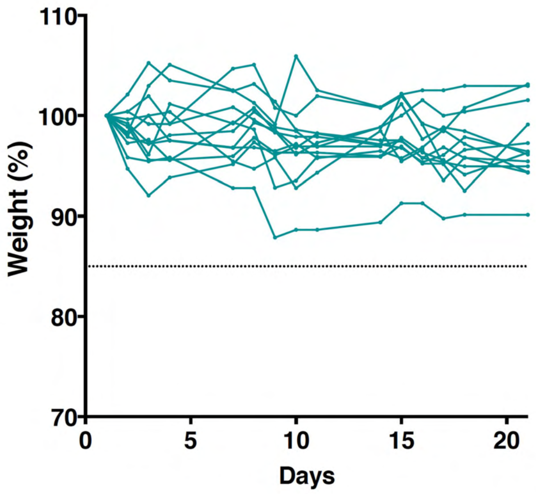


CX5461 (40 mg/kg) + Olaparib (50mg/kg) combination treatment
n=13

**Supplementary Figure 7.**

**A)** Combined CX-5461 with BMN-673 treatment does not further reduce rDNA transcription rate compared to single agent CX-5461 in OVCA cells. Cells were treated with vehicle, 1μM CX-5461, 100 nM BMN-673 or in combination for 3h. RNA was extracted and 47S rRNA precursor levels were determined using primers specific to the 5'ETS. Expression levels were normalised to NONO mRNA and expressed as fold change relative to vehicle ($n$ =3), error bars represent mean ± SEM, statistical analysis was performed using one-way ANOVA multiple comparisons, *$p$-value < 0.05, **$p$-value < 0.01, ***$p$-value < 0.001, compared to vehicle samples. NS denotes non-significant p-values. **B)** Effect of CX-5461 and olaparib treatment as described in Figure 7A&B on weight loss during treatment period.

Supplementary Table 1. TP53 mutation status assessed by high resolution melting analysis across ovarian cancer cell lines.

| Cell Line | Histology | p53 Status | Source |
|---|---|---|---|
| 2008 | Endometrioid | WT | Stephen Howell at University of California, San Diego |
| 59M | Endometrioid | c.577_592del (homo), p.His193LysfsX49 | European Collection of Cell Cultures |
| A2780 | Adenocarcinoma SEROUS | WT | European Collection of Cell Cultures |
| Caov3 | Serous | c.406C>T (homo),p.Gln136Stp | National Cancer Institute |
| CH1 | SEROUS (Adenocarcinoma) | c.948C>T (het),p.Pro316Pro | Lloyd Kelland at the Institute of Cancer Research, Sutton, UK |
| COLO704 | Adenocarcinoma | WT | Deutsche Sammlung von Mikroorganismen und Zellkuturen |
| Colo720E | Adenocarcinoma | c.1118del (het), p.Lys373ArgfsX49: c.413C>T (het),p.Ala138Val | European Collection of Cell Cultures |
| EFO21 | Serous | c.370T>C (homo),p.Cys124Arg | Deutsche Sammlung von Mikroorganismen und Zellkuturen |
| EFO27 | Mucinous | c.817C>T (het),p.Arg273Cys | Deutsche Sammlung von Mikroorganismen und Zellkuturen |
| ES2 | Serous | c.722C>T Homo),p.Ser241Phe | American Type Culture Collection |
| FUOV1 | Serous | c.535C>G (homo),p.His179Asp | Deutsche Sammlung von Mikroorganismen und Zellkuturen |
| IGROV1 | Endometriod,serous,clear cell | c.377A>G (het) ,p.Tyr126Cys | National Cancer Institute |
| JHOC5 | Clear Cell | WT | RIKEN |
| JHOC7 | Clear Cell | WT | RIKEN |
| JHOC9 | Clear Cell | WT | RIKEN |
| JHOM1 | Mucinius | c.637C>T (het),p.Arg213Stp | RIKEN |
| JHOS3 | Serous | c.783-1G>T (homo) | RIKEN |
| KURAMOCHI | Serous | c.841G>T (homo) ,p.Asp281Tyr | Health Science Research Resources Bank |
| MCAS | Mucinous | WT | Health Science Research Resources Bank |
| OAW28 | Serous | c.455del (homo), p.Pro152ArgfsX18: | European Collection of Cell Cultures |
| OAW42 | Serous | WT | European Collection of Cell Cultures |
| OV90 | Serous | c.643A>C (Homo),p.Ser215Arg | American Type Culture Collection |
| OVCA432 | Serous | WT | Dr Nuzhat Ahmed, Womens Cancer Research Centre, Royal Women's Hospital, Melbourne |
| OVCAR3 | Serous | c.743G>A (homo),p.Arg248Gln | National Cancer Institute |
| OVCAR4 | Serous | c.388C>G (homo) ,p.Leu130Val | National Cancer Institute |
| OVCAR5 | Adenocarcinoma | WT | National Cancer Institute |
| OVCAR8 | Serous | c.376-1G>A (homo) | National Cancer Institute |
| RMGI | Clear Cell | WT | Health Science Research Resources Bank |
| RMGII | Clear Cell | WT | Health Science Research Resources Bank |
| SKOV3 | Clear Cell | WT | National Cancer Institute |
| TOV112D | Endometrioid | c.524G>A (homo),p.Arg175His | American Type Culture Collection |
| TOV21G | Clear Cell | WT | American Type Culture Collection |

| Supplementary Table 2. Antibodies/Reagents | | | |
|---|---|---|---|
| **Antibody** | **Company** | **Catalogue No.** | **Application** |
| p53 (DO-1) | Santa Cruz | sc-126 | WB |
| pp53 (S15) | Cell Signaling | 9284 | WB |
| pCHK1 (S345) (133D3) | Cell Signaling | 2348 | WB |
| pCHK2 (T68) (C13C1) | Cell Signaling | 2197 | WB |
| pATM (S1981) [EP1890Y] | Abcam | ab81292 | WB |
| ATM (2C1) | GeneTex | GTX70103 | WB |
| pRPA32 S33 | Bethyl | A300-245A | WB |
| pRPA32 S4/S8 | Novus Biologicals | NB100-544 | IF |
| α-Tubulin (B-5-1-2) | Sigma | T5168 | WB |
| Actin (C4) | MP Biomedicals | 08691001 | WB |
| Goat-α-Mouse HRP | Bio-Rad | 172-1011 | WB |
| Goat-α-Rabbit HRP | Bio-Rad | 170-6515 | WB |
| Rad51 | Abcam | ab63801 | IF |
| Geminin | Abcam | ab104306 | IF |
| γH2AX (S139) [EP854(2)Y] | Abcam | ab81299 | IF |
| 53BP1 (BP13) | Merck Millipore | MAB3802 | IF |
| DNA G-quadruplex (G4) (1H6) | Merck Millipore | MABE1126 | IF |
| DNA-RNA Hybrid (S9.6) | Kerafast | ENH001 | IF |
| pATR (T1989) | GeneTex | GTX128145 | IF |
| UBF (F-9) | Santa Cruz | sc-13125 | IF |
| UBF (WT1F) | In-house | | IF |
| NPM | Abcam | Ab10530 | IF |
| Goat-α-Rabbit Alexa Fluor 488 | Invitrogen | A-11008 | IF |
| Donkey-α-Mouse Alexa Fluor 594 | Invitrogen | A-21203 | IF |
| Goat-α-Mouse Alexa Fluor 488 | Invitrogen | A-11001 | IF |

| | | | |
|---|---|---|---|
| Goat-α-Rabbit Alexa Fluor 594 | Invitrogen | A-11012 | IF |
| Vectashield | Vectorlabs | H-1000 | IF |
| Vectashield with DAPI | Vectorlabs | H-1200 | IF |
| BrdU (B44) | BD Biosciences | 347580 | FACS |
| Sheep-α-Mouse IgG FITC | MP Biomedicals | 0855520 | FACS |
| Propidium Iodide | Sigma Aldrich | P4170 | FACS |

**Supplementary Table 3**: Primer Sequences for quantitative reverse transcription real time-PCR analysis

| | Forward | Reverse |
|---|---|---|
| 47S-rRNA 5'ETS, used in Figure 1B, location (+952-1030) | GGCGGTTTGAGTGAGACGAGA | ACGTGCGCTCACCGAGAGCAG |
| 47S-rRNA 5'ETS, used in Supplementary Figure 4, location (+413-521 | GCTCTTCGATCGAGTTGGTGACG | CGGGCGGAGCGAGAAGGAC |
| Vimentin | AGAGAACTTTGCCGTTGAAGCT | GAAGGTGACGAGCCATTTCC |
| NONO | CATCAAGGAGGCTCGTGAGAAG | TGGTTGTGCAGCTCTTCCATCC |
